# Sustaining healthy long-term host-microbiome interactions in a physiologically relevant dynamic gingival tissue model

**DOI:** 10.1101/2024.01.28.577629

**Authors:** M. Adelfio, G. E. Callen, A. R. Diaz, B J. Paster, X. He, H. Hasturk, C. E. Ghezzi

## Abstract

Host-oral microbiome interactions are known to be critical in maintaining local and systemic health of the human body, although they are difficult to study in both clinical and *in vitro* applications. Despite efforts, recapitulation of gingival architecture and physiological characteristics of the periodontal niche cannot be achieved by traditional tissue engineering strategies. Here, we advanced our humanized three-dimensional gingival model by co-culturing it with a healthy patient-derived microbiomes for seven days within an oral bioreactor that mimics native salivary dynamics. Our results indicated long-term host and microbiome viability, host barrier integrity and physiological response, and preservation of healthy microbial populations and interbacterial dialogues. The model has proven useful in successfully mimicking tissue homeostasis at the interface of the periodontal niche and suitable for the introduction of immune cells. Future studies will focus on using the model as a comparator of periodontal inflammation and identifying biomarkers associated with eubiotic/dysbiotic profiles.

**Graphical Abstract:** Created partially in biorender.com.

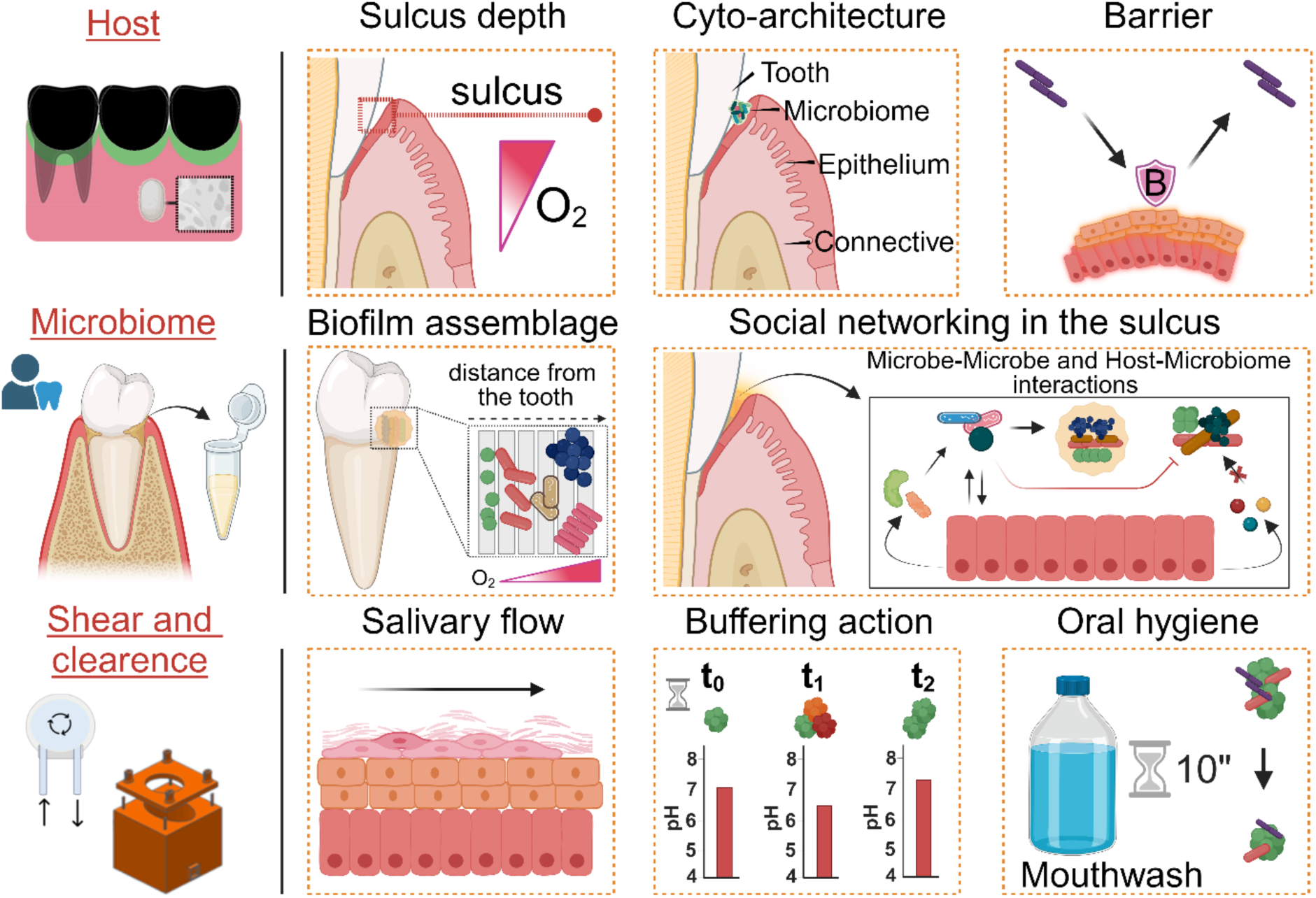

## Introduction

Although the oral cavity harbors the second largest microbiome in the human body^1^, the current understanding of host-microbiome interactions in the context of oral health is still at its infancy. Host-microbiome interactions are known to be fundamental in maintaining the local and systemic health of the human body^2^, in terms of immune regulation, barrier functions, protection against pathogen invasion or energy generation for metabolic functions^3^. Yet, all these intertwined relationships are not fully understood, nor their alteration and association with systemic diseases (*i.e.,* inflammatory bowel disease, arthritis, and Alzheimer’s disease)^4^. It has been established that the host provides commensal bacteria with a stable ecological niche in which eubiotic communities arise, assemble, and distribute according to the mouth habitat and host evolution^2,5^. However, when a biofilm shifts to a dysbiotic state, the periodontal niche can serve as a reservoir for opportunistic pathogens and lead to the onset of oral or other systemic diseases. Although advances on microbiome changes in relative abundances (from eubiosis to dysbiosis) has recently made significant progress, the conundrum that dysbiosis and persistence of dysregulated inflammation are two mutually reinforcing conditions has yet to be solved^2,5^. While the correlation between dysbiosis and disease states is clear, there is little understanding regarding the interplay between the oral tissue niche, and microbiome; thus, offering opportunities to identify predictive disease biomarkers and to develop intervention strategies to promote oral and overall health.

Given the coevolution of the microbiome with the host and their organization in multiple habitats within the depth of the gingiva^1,5^, their interactions are challenging to study in both clinical and *in vitro* applications^6^. Pre-clinical and clinical studies are limited in investigating disease trajectories, while animal models show differences in microbiome compositions, organization, and disease initiation. Current therapies for periodontal conditions remain inadequate^7^ and they are extremely costly due to routine care^6,8,9^. Yet, the application of *in vitro* research strategies using traditional tissue engineering strategies (*i.e.,* cells grown in tissue culture plastic or three-dimensional planar and static models) are still lacking several of the native human tissue nuances. The main setbacks are due to the complexity of recapitulating *in vitro* the oral host-pathogen niche features (*i.e.,* oxygen and pH gradients) and microenvironmental stimuli (*i.e.,* salivary flow, drinking, chewing) that ultimately dictates the continuously evolving relationship between the host and microbial populations to allow mutual coexistence within the oral environment^1,10^. Despite the recent research efforts^6,11,12^, a physiologically relevant model, encompassing all of these features, is still representing an unmet challenge in the field of oral tissue modeling. Underrepresentation of salivary composition and shear forces, fundamental in regulating tissue homeostasis through the maintenance of the ecosystem stability and the mucosal turnover^2^, strongly contributed to failed attempts in co-culturing mammalian cells and patient-derived microbiomes^13^. Difficulties also arise from not adequately balancing microbiome richness and abundance due to the lack of a native oral tissue architecture, which ultimately led to the use of bacterial lines grown on three-dimensional oral-mucosa equivalent^12–16^. The use of bacterial cell lines in combination with mammalian cells together with the lack of native tissue structural features resulted in significant limitations: (*i*) disregarding the concept of microbial ecology, self-assembly, and host-microbial coexistence on non-shedding surfaces^1,13,17,18^; (*ii*) overly reducing the role of the host in modulating biofilm composition through salivary flow, antimicrobial peptide release factors and anatomy-determined oxygen and pH gradients^1^; (*iii*) limited opportunities to identify disease biomarkers to advance intervention strategies, as they do not resemble *in vivo* conditions. Despite recent progress, challenges and prospective failure points persist; thus, there is still a compelling need for improved current *in vitro* technologies.

Currently, there are no experimental *in vitro* models able to mimic the structural, physical, and metabolic conditions present in the oral cavity to support the long-term investigation of host-pathogens interactions. We have previously developed a three-dimensional humanized gingival tissue model based on natural proteins to recreate the anatomical architecture of an adult human lower-jaw gingiva. We have found that the ability to replicate the anatomical tissue architecture is crucial for the establishment of the physiological oxygen gradients within the periodontal pocket to support the long-term culture of microbial populations^6^. The model sustained interactions between human primary gingival cells and patient-derived subgingival plaque-derived microbiomes for up to 24 hours, while preserving microbiome richness and diversity. To meet the challenge of long-term studies and align with experimental criteria dictated by clinical studies^7^, the model was integrated into a bioreactor to mimic salivary flow composition, shear stress and buffering action^10^ and oral hygiene was simulated by rinsing with a commercial mouthwash. In the present work, we described an *in vivo-like* environment to investigate long term interactions between patient-derived gingival tissue and oral microbiome, displaying the establishment of symbiotic relationships through various biological, molecular, and functional redouts. This oral cavity model showed pro-inflammatory responses to microbial challenge (day 1 post-inoculation), sustained physiological anti-microbial activity, cytokine responses similar to pre-clinical *in vivo* features, and the maintenance of a viable and eubiotic microbiome. In conclusion, we have developed a model able to recreate the *in vivo* homeostatic balance between host and microbiome. Future studies will focus on implementing this *in vitro* model system as a platform to investigate the correlation between dysbiosis and inflammation and the efficacy of inflammation resolution strategies to regress severe periodontal disease in alignment with clinical studies.

## Materials and Methods

### Anatomical model preparation

#### Aqueous silk solution

Silk solution was prepared from *B. morii* silkworm cocoons (Tajima Shoji Co., Yokohama, Japan) according to the established protocol^19^. 5 gr of silk cocoons were cut into small pieces and degummed for 30 minutes to remove sericin protein in 2L of deionized (D.I.) water, containing 0.02 M sodium carbonate (Thermo-Fisher Scientific, Waltham, MA). The resulting fibers were rinsed three times (20 minutes/rinse) in D.I. water and dry overnight before being dissolved in in 9.3 M lithium bromide (Sigma-Aldrich, St. Louis, MO) solution for 2 hours at 60°C. Subsequently, the obtained solution was dialyzed against D.I. water for three days in a cellulose dialysis tube (3.5 kD MWCO, Spectrum Labs Inc, Rancho Dominguez, CA) for a total of six D.I. water changes and lastly collected and centrifuged two times at 9000 RMP for 20 minutes at 4°C. Aqueous silk solution concentration was assessed by weighing a dried sample of known volume and diluted to 4% w/v in D.I. water.

#### Silk sponge scaffolds with gingival anatomical architecture

Gingival anatomical scaffolds were made by replica molding, as previously described^10^. Briefly, silk solution (4 wt/%, 4̴ mL/mold) was pipetted into polydimethylsiloxane (PDMS) mold replicating a three gum-tooth unit (lower jaw), degassed, and then freeze-dried (Labconco, MO) to induce formations of a porous structure. The resultant scaffolds were autoclaved to induce insolubility in water via changes in β-sheet crystallinity structure. For cell culture purposes, silk sponges were autoclaved a second time.

### Cell culture and artificial saliva preparation

#### Oral primary gingival cells

Oral stromal and keratinocyte cells were purchased from Lifeline Technologies (Lifeline Cell Technologies, Frederick, MD). Cells were grown at 37°C and 5% CO_2_, cultured up to passage 6 as per manufactory’s instructions and Lifeline Trypsin/EDTA 0.05/0.02% and Lifeline Trypsin Neutralization Solution were used for cellular detachment. Oral stromal cells (hGFCs) (FC-0095) were cultured in FibroLife serum-free media (LL-0001) supplemented with FibroLife serum-free LifeFactors containing: (HLL-Human serum Albumin (500 µg/mL), Lecithin (0.6 µg/mL), Linoleic Acid (0.6 µM)), Epidermal Growth Factor/Transforming Growth Factor-β1 – rhEGF/TGFβ1 (5 ng/mL, 30 pg/mL), Fibroblast Growth Factor-rhFGF (5ng/mL), rhInsulin (5 µg/mL), L-Glutamine (7.5 mM), Ascorbic acid (50 µg/mL), Hydrocortisone Hemisuccinate (1 µg/mL) and Gentamicin and Amphotericin B (30 µg/mL, 15 ng/mL). Oral Keratinocyte cells (hGECs) (FC-0094) were cultured in DermaLife basal medium (LL-0007) supplemented with DermaLife K LifeFactors kit containing: rhTGF-α (0.5 ng/mL), rhInsulin (5 µg/mL), L-Glutamine (6 mM), Extract PTM (0.4%), Epinephrine (1 µM), Apo-Transferrin (5 µg/mL), Hydrocortisone Hemisuccinate (100 ng/mL) and Gentamicin and Amphotericin B (30 µg/mL,15 ng/mL).

#### Artificial saliva preparation

Artificial saliva was prepared as previously described^10^. Briefly, *Xanthan gum* (x. gum) (0.05 wt%) (Sigma-Aldrich, St. Louis, MO) was dissolved in optimized co-culture media composed of FibroLife serum-free media and DermaLife Calcium-Free basal media (LL-0029)^20^ in a 3:1 ratio and supplemented with fibroblast and keratinocyte growth^21^. Subsequently, the artificial saliva was supplemented with the major human salivary ionic components adapted by Sarkar and colleagues^22^: sodium chloride (1.594 mg/mL), potassium phosphate (0.636 mg/mL), potassium chloride (0.202 mg/mL) purchased from Thermo-Fisher Scientific (Waltham, MA) and ammonium nitrate (0.328 mg/mL), potassium citrate (0.308 mg/mL), uric acid sodium salt (0.021 mg/mL), urea (0.198 mg/mL), and lactic acid sodium salt (0.146 mg/mL), purchased from Sigma-Aldrich (St. Louis, MO). Artificial saliva (pH 7.4) was filtered using a 100 µm strainer and a vacuum filtration system (0.45 µm) (VWR, Philadelphia).

### Anatomical inoculated gingival tissue model: native stimulation

#### Bioreactor fabrication

The bioreactor was fabricated and machined at the Lawrence Lin MarkerSpace-UMass Lowell, as previously described^10^. The bioreactor set-up was composed of a chamber equipped with a lid made in Derlin® acetal resin (McMaster-Carr, Elmhurst, IL), a peristaltic pump (EZO-PMPTM-Atlas Scientific, Long Island City, NY) and an Arduino set-up (board and breadboard). The chamber was sealed with pin screws, flat washers, wing nuts and an O-ring (McMaster-Carr, Elmhurst, IL) and connected to a peristaltic pump via tubing and connectors (McMaster-Carr, Robbinsville, NJ; United States Plastic Corporation, Lima, OH; Masterflex, Vernon Hills, IL and Atlas Scientific, Long Island City, NY). All components were autoclaved at 121°C for 20 minutes prior to the culture. Each pump functioned through an Arduino set up (board and breadboard) and a code written in Arduino (https://atlas-scientific.com/peristaltic/ezo-pmp/).

#### Cellular gingival scaffold preparation

Anatomical scaffolds (**Figure 1**) were prepared as previously described^10^. Briefly, scaffolds were pre-processed by gently squeezing out of the water content before soaking them in Phosphate Buffer Saline (PBS) (1x) and then drying them on sterile gauze pads. During the drying process, hGFCs were gently detached and incorporated (200,000 cells/mL) in neutralized rat tail collagen type I solution (First Link, UK) before being seeded onto the scaffold (closed porosity). Collagen solution was made at 4°C by combining acidic collagen and Dulbecco’s Modified Eagle Medium (DMEM, 10x) in a 4:1 ratio, and neutralized with sodium hydroxide (10 M) (Sigma-Aldrich, St. Louis, MO). Cellularized scaffolds were incubated for 30 minutes at 37°C to induce collagen polymerization. Subsequently, hGECs were gently detached and seeded, twice, in the periodontal pocket (open porosity) at 50,000 cells/cm^2^, with a two-hours interval between seedings to support cell adhesion. Gingival scaffolds were cultured in co-culture media for two days and then in artificial saliva for the entire duration of the experiment (2 weeks). On day 5 post-seeding, scaffolds were cultured in air-liquid-interface (ALI) conditions and dental resin teeth^6^ were inserted into the pockets to complete the anatomical structure. Artificial saliva was replaced three times a week for the first seven days and then every day after the subgingival plaque microbiome inoculum (**Figure 1**).

**Figure 1.**
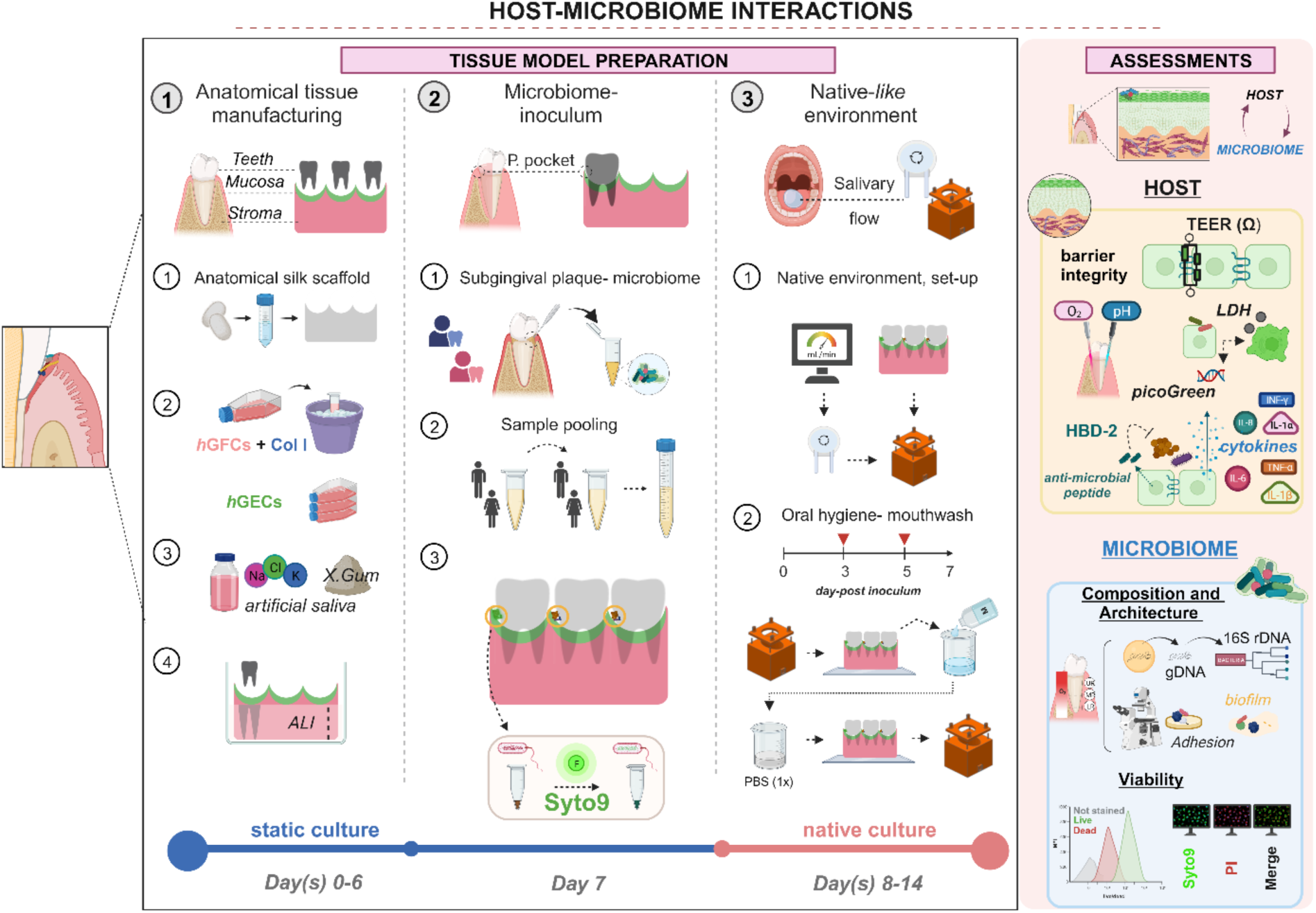
*In vivo* mimicking long-term host-microbiome interactions: experimental workflow. Three main steps: 1) *Anatomical tissue manufacturing:* silk-based constructs with anatomical architecture were populated with primary keratinocytes (hGECs) and human gingival stromal cells (hGFCs) delivered in type I collagen and cultured in artificial saliva until scaffold maturation. 2) *Microbiome inoculum:* patient-derived subgingival plaque microbiomes (N_patients_=8) were pooled, inoculated into periodontal pockets in the scaffolds and cultured under static conditions for 16h; microbiome was pre-stained with Syto9 dye for viability tracking. 3) *Native-like environment:* before starting the native conditions, the bioreactor setup was calibrated to ensure the correct flow rate. Host-microbiome interactions were established for seven days, and on the third and fifth days after inoculation, oral hygiene was simulated by rinsing with a commercial mouthwash. Host-microbiome responses were assessed by molecular and biological readouts. Figure created partially with biorender.com.

#### Oral subgingival plaque microbiome: inoculum

Human subgingival plaque microbiome samples were collected from healthy donors (N=8 with 10 sampling sites/donor) at the Center for Clinical and Translational Research at ADA Forsyth Institute, Cambridge (MA). Before seeding, plaque samples were pooled and split for the following analyses: genomic DNA extraction for 16S rDNA analysis (*h*-plaque 0), Syto9 dye incubation (pre-seeding), flow cytometry analysis (*h*-plaque 0) and model seeding. Plaque samples were then resuspended in artificial saliva and inoculated into the gingival sulcus of each scaffold in 10 µL aliquots (5 µL per pocket side) on the 7^th^ day post-seeding (day 0 post-inoculum). Inoculated scaffolds (N=5) were maintained in static conditions at ALI for 16 hours to facilitate bacterial attachment and subsequently cultured in native regime (1 mL/min). Inoculated oral tissue models (OTMs) were cultured for seven days. All methods were performed in compliance with the approved protocol of ADA Forsyth Institutional Review Board (Protocol No. FIRB# 18-06) and the University of Massachusetts Lowell (Protocol No. IRB - 20-090). In addition, subjects signed the FIRB-approved informed consent prior to sampling.

#### Host-microbiome interactions: native regime

To induce and replicate salivary shear stimulation, OTMs were transferred into the bioreactor chamber and cultured in native regime (1 mL/min) at 37°C and 5% CO_2_. Before proceeding with the native regime and each day for media (artificial saliva) change, the peristaltic pumps were calibrated to ensure accurate flow rate. Host-microbiome early-response was performed for three days (day 3 post-inoculum), in which host and microbiome’s interactions were evaluated both in static (N=3) and native (N=3) conditions. Artificial saliva was replaced every day and aliquots from every chamber reservoir and from each periodontal pockets (N=3 per OTM) were taken for further analyses, such as, pH measurements, Lactate Dehydrogenase (LDH) assay, enzyme-linked immunosorbent assay (ELISA), and Milliplex cytokines panel.

#### Host-microbiome interactions: mimicking oral hygiene

To mimic daily oral hygiene routines, we performed oral rinses using commercial mouthwashes. We first investigated cytotoxicity of three mouthwashes (Listerine®, Colgate® and Crest®, in %: 0,1,2,5,8) in PBS (1x) on hGFCs via live/dead assay (Thermo-Fisher Scientific, Waltham, MA). hGFCs were seeded on planar silk sponge scaffolds at the density of 228.8 cells/cm^2^ and grown for two days. In order to mimic human clinical conditions, cells were incubated for 30 seconds with the mouthwash, rinsed with PBS (1x), and then incubated for 15 minutes at room temperature with live/dead dyes. Live/Dead analysis was assessed using both a microplate reader (Emission/Excitation (nm): Live (494/517) and Dead (528/617), SpectraMax M2 with SoftMax Pro7 software, Molecular device, San Jose, CA) and a fluorescence microscope (Life technologies EVOS FL). After a preliminary investigation with hGFCs, we down selected and exposed hGECs (1,287 cells/cm^2^) to only Listerine®. Oral rinses in the anatomical scaffolds were performed on day 3 and day 5 post-inoculation using Listerine® Antiseptic mouthwash original (brown color) diluted in PBS (1x) to 1% and filtered using a vacuum filtration system (0.2 µm) (VWR, Philadelphia). Briefly, scaffolds were inserted into a sterilized beaker containing enough mouthwash solution to reach ALI conditions on the scaffold, and rinsed for 10 seconds, before being transferred to another sterilized beaker containing PBS (1x).

### Host -microbiome interactions - Host

#### Host viability assessments: lactate dehydrogenase (LDH) and picoGreen

Longitudinal monitoring of host viability within the OTM was carried out via LDH assay to indirectly quantify cytotoxicity and picoGreen assay for DNA quantification. LDH assay was conducted according to the manufacturer’s instructions (Sigma-Aldrich, St. Louis, MO). 500 µL of spent saliva were stored at - 80°C, then diluted 1:5 in LDH assay buffer and obtained readings were interpolated using the standard curve. Mammalian host cells were isolated microbial cells using Percoll^TM^ (Cytiva, Marlborough, MA) density gradient^23^, and then quantified using Quant-iT picoGreen dsDNA assay kit (Thermo-Fisher Scientific, Waltham, MA), according to manufacturer instructions. Briefly, epithelium was isolated from the scaffold via nasal swabs^24^, vortexed briefly in PBS (1x) and centrifuged at 500 xg to remove some of the bacterial cells. The pellet was then resuspended in 1 mL of isolation medium composed of 50 µL of DermaLife basal medium supplemented with L-Glutamine (6 mM) (Lifeline Cell Technologies, Frederick, MD), 10% Fetal Bovine Serum and 2.5 nM of L-Cysteine (Thermo-Fisher Scientific, Waltham, MA) plus 950 µL of 50% v/v Percoll^TM^ in PBS (1x). The cells were centrifuged at 11,000 xg for 15 minutes, then the first 500 µL containing only the epithelial cells were mixed with PBS (1x) and centrifuged again at 500 xg for 5 minutes; the remaining 500 µL containing the microbiome were processed for flow cytometry (section: *Host-microbiome interactions – microbiome: flow cytometry*). To remove any bacterial contamination, the pellet was resuspended in PBS (1x) and filtered through a 12-µm hydrophilic polycarbonate Nucleopore (Whatman) filter; finally, the membrane was briefly vortexed, and the recovered epithelial cells were gently resuspended in 0.5 mL of 0.05 % Triton-X (Sigma-Aldrich, St. Louis, MO) and stored at -80°C before proceeding with a standard picoGreen protocol.

#### Oral barrier: TransEpithelial Electrical Resistance (TEER) and confocal laser scanning microscopy (CLSM)

Oral epithelial barrier integrity was assessed using a voltohmmeter (EVOM^2^) and the associated cell culture cup chambers-6mm (ENDOHM-6G) (World Precision Instrument, Sarasota, FL). OTMs on day 3 or day 7 post-inoculum (day 10/14 post-seeding) were punched into 7 mm discs using a biopsy punch (VWR, Philadelphia) to fit into a Transwell insert (#29442-082, VWR, Philadelphia) and TEER values were recorded. TEER values (equation 1) were reported as Ω/mm and thickness of the specimen was measured using a caliper. Blank was assessed by recording TEER in collagen-filled acellular anatomical scaffolds (N=5) in artificial saliva.

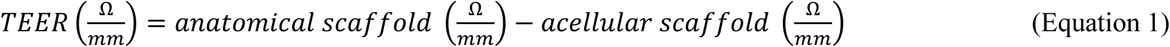

To assess epithelium stratification and differentiation after microbiome interactions, OTMs on day 3 post-inoculum were processed for immunohistochemistry^10^. Briefly, the scaffolds were rinsed in PBS (1x), and fixed in Paraformaldehyde (PFA) (4%) (Thermo-Fisher Scientific, Waltham, MA) for 2 hours at room temperature (RT). Scaffolds were then permeabilized in cold Acetone (100%) (Sigma-Aldrich, St. Louis, MO) for 10 minutes at -20°C and blocked in Bovine Serum Albumin (BSA, 1%) (Sigma-Aldrich, St. Louis, MO) in PBS-Triton (PBS-T, 0.5%) for 1 hour at RT. Incubation with the primary antibody was carried out at 4°C overnight in a PBS (1x) solution containing anti-Ki67 (Abcam, MA, 1:100), BSA (1%), horse serum (10%) (Fisher-Scientific, Hampton, NH) and PBS-T (0.5%). Secondary staining was performed for 2 hours at RT in a PBS (1x) solution containing secondary antibody-TRIC labeled (Abcam, MA, 1:1000), BSA (1%) and PBS-T (0.5%) followed by counterstaining with 4’,6-diamidino-2-phenylindole (DAPI) in PBS (1x) (Sigma-Aldrich, St. Louis, MO, 1:1000) for 20 minutes at RT. Scaffolds were imaged with a confocal laser scanning microscope (CLSM) Leica SP8 (Leica, Germany).

#### Oral mucosa responses: anti-microbial peptides release

Analysis of anti-microbial peptide human β-defensin 2 (PeproTech, Cranbury, NJ - 900-K172) was performed in OTMs via ELISA. Spent-artificial saliva samples (100 µL) were collected from each periodontal pocket on day 0 (pre-inoculum) and on day 1, 2, 3, 5 and 7 post-inoculum and stored at -80°C until analyzed. Due to bacterial and debris interferences with ELISA plates, samples were briefly spun down before being frozen. Human β-defensin 2 detection was performed according to manufacturers’ instructions and samples were diluted 1:4 in reagent diluent, as indicated by the manufacturer. Additional reagents were purchased separately: microplates (Corning, NY), Tween-20, Bovine Serum Albumin and ABST liquid substrate solution (Sigma-Aldrich, St. Louis, MO) and PBS (10x) (Thermo-Fisher Scientific, Waltham, MA). Optical Density (OD) was determined using a microplate reader (SpectraMax M2 with SoftMax Pro7 software, Molecular device, San Jose, CA) set at 405 nm and wavelength correction was applied by subtracting readings at 650 nm from 405 nm. All OD values were then interpolated based on a standard curve.

#### Oral anti and pro-inflammatory responses: milliplex assay

Epithelium exudates were collected from each periodontal pocket in the OTMs on day 0 1, 3 and 7 post-inoculums. Simultaneous analysis of multiple cytokine and chemokine biomarkers for inflammation (Granulocyte-macrophage colony stimulating factor (GM-CSF), Interferon gamma (IFN-γ), Interleukin 10 (IL-10), Interleukin 1 Receptor Antagonist (IL-1RA), Interleukin 1 alpha (IL-1α), Interleukin 1 beta (IL-1β), Interleukin 4 (IL-4), Interleukin 6 (IL-6), Interleukin 8 (IL-8), Monocyte Chemoattractant Protein 1 (MCP-1), Tumor Necrosis factor alpha (TNF-α)) was carried out with human cytokine/chemokine magnetic bead panel (MilliPlex MAP, Millipore, Sigma). Median fluorescence intensity (MFI) in each sample was acquired using the Luminex 200 instrument (Luminex, Texas) and the associated program (xPonent); the raw data were then interpolated by using a standard curve by means of Belysa® Immunoassay curve fitting software (Merck Millipore, Burlington, MA). Resulting protein concentration (pg) were then normalized for the respective cellular density obtained by Percoll^TM^ (section: Host viability assessments: lactate dehydrogenase (LDH) and picoGreen).

#### Localized gradients at the periodontal niche: pH and Oxygen

To monitor changes in local environment conditions due to host and microbiome metabolic interactions, pH and oxygen were measured in the periodontal pocket. Oxygen was recorded using the PreSens’s Profiling Oxygen Microsensor PM-PSt7 with the Microx 4 fiber optic oxygen transmitter (PreSens, Regensburg, Germany) inserted from the upper to the lower regions of the pocket by means of Narishige’s M-152 Three-Axis Direct-Drive Coarse Micromanipulator and GJ-1 Magnetic Stand with steel base plate (Narishige, Amityville, New York). pH values (100 µL/pocket) were recorded by using the Orion™ 9810BN Micro pH Electrode (Thermo-Fisher Scientific, Waltham, Massachusetts) and the FiveEasy™ pH/mV Meter (Mettler-Toledo, Columbus, OH).

### Host-microbiome interactions – Subgingival plaque microbiome

#### Microbiome viability assessment: confocal laser scanner microscopy (CLSM)

To characterize microbial viability and distribution within the model, pooled subgingival microbiome samples were pre-stained with Syto9 (Thermo-Fisher Scientific, Waltham, MA) prior to inoculum. Briefly, pooled subgingival plaque microbiome was incubated for 20 minutes in a solution containing Syto9 (1:250) in PBS (1x) at RT to ensure Syto9 binding to microbial nucleic acids, subsequently inoculated, as previously described. On day 7 post-inoculum, pockets were incubated with Propidium Iodide (PI) (30 µM) (Thermo-Fisher Scientific, Waltham, MA) in PBS (1x) for 20 minutes at RT to stain for dead microbial cells. OTMs were then washed in PBS (1x) and imaged with a CLSM Leica SP8 (Leica, Germany) to characterize microbiome viability and distribution.

#### Microbiome viability assessment: flow cytometry

To monitor microbial viability over time, live/dead assay was performed via flow cytometry on day 0, 3 and 7 post-inoculums. Microbial samples (500 µL) on day 3, 7 post-inoculums were isolated from the OTMs via Percoll^TM^ density gradient (see section: Host viability assessments: lactate dehydrogenase (LDH) and picoGreen) and processed for live/dead assay. Pooled subgingival microbial samples were resuspended in 0.85% sodium chloride buffer (Thermo-Fisher Scientific, Waltham, MA) and grouped in live (stained only with Syto9, Syto9+PI and not stained) and dead controls (stained only with PI or Syto9+PI). Negative controls were obtained by incubation in 70% isopropanol (Thermo-Fisher Scientific, Waltham, MA) for 30 minutes at RT with subsequent vortexing. Samples were then incubated for 15 minutes at RT with Syto9 and/or PI (Thermo-Fisher Scientific, Waltham, Massachusetts) at 20 and 60 µM final concentration in 0.85% sodium chloride buffer, respectively. After incubation, samples were fixed in PFA 2% in PBS (1x) (Thermo-Fisher Scientific, Waltham, Massachusetts) before being analyzed for flow cytometry using Amnis® FlowSight® Imaging Flow Cytometer (Millipore Sigma, Burlington, MA). 10,000 events/sample were collected, and the specific subpopulation of single cell was selected through a customized gating strategy. Before acquisition, a compensation matrix was performed using subgingival microbiome samples^25^. Subsequently, the data were analyzed with IDEAS software (Amnis Corporation, Seattle, WA). Briefly, the Gradient RMS function on the free-field channel (Ch01) was used to identify and eliminate in-focus events; then, a scatter plot of area *vs* aspect ratio of Ch01 was performed to identify the desired population and exclude debris. The selected population was then gated for Syto9 (Ch02) *vs* PI (Ch04) to analyze the percentage of cells positive for each dye (**Supplementary Figure 7a**).

#### Microbiome spatial organization within the OTM: scanning electron microscope (SEM)

To characterize microbiome spatial organization, OTMs were rinsed with PBS (1x) and then fixed in PFA 4% (Thermo-Fisher Scientific, Waltham, MA) in PBS (1x) for 2h at RT. OTMs were cut longitudinally and sliced transversally to image the inside portion of the pocket. OTMs were rinsed in D.I. water and then dehydrated through graded ethanol series (in %: 50, 70, 80, 90, 95 and 100) (Thermo-Fisher Scientific, Waltham, MA) before proceeding with Critical Point Dryer (CPD) (Tousimis Samdri, Rockville, MD). OTMs were then coated with Au/Pd particles (Cressington 108) (Cressington Scientific Instruments, UK) and imaged with a Phenom XL G2 Desktop (Thermo-Fisher Scientific, Waltham, MA) equipped with an EDS sensor at 10 kV accelerating voltage and low vacuum (60 Pa). OTMs on day 3 post-inoculum were coated with gold particles (Denton Vacuum Desk IV Sputter Coater, Moorestown, NJ) and subsequently imaged with the JEOL 6390 SEM (Peabody, MA) using a 5-10 kV accelerating voltage.

#### Assessment of microbiome taxonomy: gDNA isolation

In order to amplify only the gDNA from live bacteria, we implemented a method to intercalate the DNA from dead bacteria with propidium monoazide (PMA) so that it does not get amplified during PCR. We performed an initial pilot experiment to validate this experimental procedure. Human subgingival plaque microbiome samples (N=4) from healthy donors were pooled and centrifuged for 3 minutes at 10,000 xg. The pellet was resuspended in 0.4 mL of PBS (1x) and divided equally into four tubes (Live Microbiome PMA-not treated, Live Microbiome PMA-treated, Heat-Killed Microbiome PMA-not treated, Heat-Killed Microbiome PMA-treated). To induce microbial death, samples were heat-killed at 90°C for 5 minutes. Samples were subsequently incubated with PMA 50 µM for 10 minutes at RT on a shaker covered by aluminum foil, before being treated with a blue LED photolysis device (Biotium, San Francisco) for 10 minutes, adopted for photoactivation of PMA. Lastly, microbial samples were centrifuged for 10 minutes at 5,000 xg and incubated overnight with Ready-Lyse lysozyme solution in TE buffer at 37°C. gDNA isolation was performed using the Epicentre MasterPure Gram Positive DNA Purification Kit (Lucigen, Middleton, WI, USA) according to manufacturer’s instructions and total gDNA was quantified using a nano-drop (Thermo-Fisher Scientific Waltham, MA). To verify the photoactivation of PMA in not amplifying gDNA from dead microbial cells, an endpoint PCR (Biorad, Hercules, California) was performed using custom primers for 16S rRNA gene (V1-V3 regions)^26^ (27F:AGAGTTTGATC**M**TGGCTCAG; 518R:GTATTACCGCGGCTGCTGG) (Thermo-Fisher Scientific Waltham, MA). Master mix (Thermo-Fisher Scientific Waltham, MA) components for a final volume of 50 µL were: Mastermix (1x, 25 uL), Forward and Reverse primers (0.5 µM, 2.5 µL), gDNA (20 ng, 1µL) and nuclease free water (up to volume). Based on primers annealing temperatures, PCR conditions were: initial denaturation at 94°C for 2 minutes (1 cycle), denaturation at 94°C for 1 minute, annealing at 94°C for 2 minutes and extension at 72°C for 2 minutes repeated for 20 cycles; final extension at 72°C for 5 minutes. PCR products were mixed 1:6 with a loading dye and run on an agarose gel (1%) in Tris-acetate-EDTA buffer supplemented with 0.2 µG/mL of Ethidium Bromide (Biorad, Hercules, California). DNA was visualized under a short-wavelength UV light to confirm the lack of amplification for Heat-Killed Microbiome PMA-treated in comparison to the Heat-Killed Microbiome control (**Supplementary Figure 2**).

Upon validation of PMA treatment to selectively amplified only live microbial organisms, on day 0 and 7 post-inoculum, microbial genomic DNA (gDNA) was isolated for 16S rDNA sequencing. For gDNA isolation on day 0 (*h*-plaque 0), 10 µL of the pooled subgingival plaque microbiome (N=5) were dispensed in 1.5 mL tubes containing 390 µL of PBS (1x) before preceding to the blue LED photolysis treatment, as described above. For samples collected on day 7 post-inoculum, one pocket per sponge (N=5) was divided into three regions-upper, medium, and lower-, using a razor blade, according to previously identified oxygen content (**Figure 1** and **Supplementary Figure 6**)^6^. Each region was manually disrupted using dissection scissors (VWR, Philadelphia) and resuspended in 400 µL of PBS (1x). Samples on day 7 post-inoculum were incubated with PMA before being treated with a blue LED photolysis device (Biotium, San Francisco), as described above. Photoactivation of PMA was performed for 15 minutes (*h*-plaque 0/day 0) or 30 minutes (day 7 post-inoculum); longer exposure was performed according to the manufacturing instructions due to the greater complexity of the microbial samples. After treatment, samples were centrifuged and resuspended in TE buffer and incubated overnight at 37°C after addition of the Ready-Lyse lysozyme solution (Lucigen, Middleton, WI, USA). gDNA extraction (Lucigen, Middleton, WI) was performed according to manufacturer’s instructions and gDNA content was quantified using a nanodrop (Thermo-Fisher Scientific, Waltham, MA). 300 ng of gDNA/sample were processed for sequencing analysis, according to Zymo Research’s guidelines (Irvine, CA).

#### Assessment of microbiome taxonomy: library preparation and 16S rDNA sequencing

After gDNA isolation, samples were shipped to Zymo Research (Irvine, CA) for library preparation and 16S rDNA sequencing. gDNA samples were prepared for targeted sequencing with the Quick-16S™ NGS Library Prep Kit (Zymo Research, Irvine, CA). Primers were custom designed by Zymo Research to ensure best coverage of the 16S gene and high sensitivity - Quick-16S™ Primer Set V1-V3 (Zymo Research, Irvine, CA). The final PCR products were quantified with qPCR fluorescence readings and pooled together based on equal molarity. The final pooled library was cleaned up with the Select-a-Size DNA Clean & Concentrator™ (Zymo Research, Irvine, CA), then quantified with TapeStation® (Agilent Technologies, Santa Clara, CA) and Qubit® (Thermo Fisher Scientific, Waltham, WA). Final libraries were sequenced on Illumina® MiSeq™ with a V3 reagent kit (600 cycles) and sequencing was performed with 10% PhiX spike-in.

#### Assessment of microbiome taxonomy: data analysis

Comprehensive data analyses and interpretation of the reads were performed at ADA Forsyth Oral Microbiome Core (FOMC, Cambridge, MA). For the analysis, a DADA2 pipeline was used^27^ to model and correct errors in the amplicon reads generated through Illumina-sequencings. DADA2 pipeline included five steps: 1) read trimmering based on sequence quality, 2) learn the error rates, 3) infer amplicon sequence variants (ASVs), 4) merge paired reads and 5) remove chimera. After DADA2 read processing, the generated ASVs were used for microbial profile comparison and taxonomy assessments. To identify oral, or not, microbial species, each sequence was screened though several databases (Expanded Human Oral Microbiome Database-eHOMD vs 15.2^28^, HOMD 16S rRNA RefSeq, Mouse Oral Microbiome Database-MOMD^29^, Green Gene Gold-GC and the NCBI 16S rRNA reference sequence set. The final OTUs were assigned based on algorithm^30^ with parameters of reads with ≥ 98% sequence identity to the matched reference and alignment length ≥ 90%. If a read matched with reference sequences representing more than one species with equal percent identity and alignment length, it was subject to chimera checking with USEARCH program version v8.1.1861^31^. Lastly, this analysis generated a read count for each sample toassess relative abundance 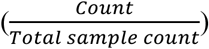 at different taxonomy level (Domain to Species).

Quantification of shared and unique Operational Taxonomic Units (OTU) between day 0 (*h*-plaque 0) and day 3 or day 7 post-inoculum was achieved and illustrated though a Venn diagram. Venn diagram analysis was performed by binarizing data from relative species abundance (0: absent; 1:present) and counting species among all possible combinations (shared or unique) in the experimental groups. Species missing in all analyzed groups for a given comparison were excluded from the count.

To quantitively describe microbiome diversity (richness, evenness, and similarity) between day 0 (*h*-plaque 0) and day 3 or day 7 post-inoculum, α and β-diversity indices were analyzed. α-diversity indices (Observed features, Shannon, and Simpson) were calculated to assess the initial (*h*-plaque 0) richness and evenness with respect to the microbial population after seven days in the OTMs (native day 7 post-inoculum). α-diversity statistical differences among different comparison groups were assessed with the Kruskal Wallis H-test (alpha-group-significance function QIIME 2 “diversity” package). β-diversity was used to assess similarity, or dissimilarity, between the initial inoculate (*h*-plaque 0) and the microbial population after seven days in the anatomical models (native day 7 post-inoculum). Count data were transformed into centered log ratio (CLR) and data were plotted using Principal Component Analysis (PCoA) using Euclidean distances (Aitchison distance). To test the distance between the two groups (*h*-plaque0 and microbiome in the OTM-native day 7 post-inoculum), a PERMANOVA (permutational multivariate analysis of variance) (beta-group-significance function QIIME 2 “diversity” package) test was performed.

To compare the proportion of different species in a sample and consider the compositional nature of the data, we performed differential abundance analysis. Differential abundance (DA) analysis of microbial taxa was assessed using the statistical tool ANCOM-BC2 (Analysis of Compositions of Microbiomes with Bias Correction)^32^, which 1) considers the compositional nature of the data, and 2) corrects the bias (*i.e.,* differences in sequencing depth) introduced by the sequencing process. The obtained DAs were then modeled using regression models for pairwise comparisons with regard of mixed directional false discover rates (mdFDR)^32–34^(**Supplementary material_16S rDNA_ANCOM-BC).** DAs were also analyzed using LEfSe (Linear Discriminant Analysis Effect Size). LEfSE analysis consisted of 2-stage statistical analysis: 1) a rank-based Kruskal-Wallis sum-rank test, to detect the significant DA between the two groups (h-plaque 0 vs native day 7), and 2) an unpaired Wilcoxon rank-sum test, to assess the biological consistency^35^. Lastly, significant DAs were used to calculate the effect sizes via linear discriminant analysis (LDA) scores.

To evaluate the social structure of the microbiome, a similarity matrix was generated (https://software.broadinstitute.org/morpheus/) from species relative abundance values based on Spearman rank correlation and imported in a custom-made MATLAB script. Briefly, community_louvain.mat script^36^ have been used (inputs: similarity matrix; gamma=1 (classic modularity); ‘negative_sym’ for symmetric treatments of negative weights) to compute the modularity value and assign each species to a module. The resulting module list was used to re-arrange the order of the species in the heatmap to highlight the modularity structure.

To identify trends based on the temporal trajectories of analytes, agglomerative hierarchical clustering with linkage analysis was applied to the cytokine panel using the average method and correlation distance metric. Visual inspection of both the heatmap and dendrogram was used to determine an appropriate cutoff threshold. Moreover, hierarchical clustering of relative abundance values of genera was generated by importing the values into the open-source software (https://software.broadinstitute.org/morpheus/). Hierarchical clustering was based on 1-Spearman rank correlation, as metric, and average, as linkage method.

Cohen’ s effect size between comparisons (**Supplementary Table 1**) was computed from α-diversity indices values (mean and standard deviation) by using the open-source calculator (https://lbecker.uccs.edu/).

### Statistical analysis

Statistical significance was calculated using OriginPro 2023b (OriginLab Corporation, Northampton, MA, USA). Statistical significance among normally distributed data was tested using two-sample Student’s T-test, Paired Samples t-test, One-Way ANOVA, One-Way or Two-way repeated measures ANOVA with Bonferroni and Dunnet’s post-hoc, p-values<0.05. Analysis of sequencing data was performed by ADA Forsyth Oral Microbiome Core (FOMC) at ADA Forsyth institute (Cambridge, MA) using: Kruskal Wallis H-test and PERMANOVA (QIIME 2 “diversity” package), and Kruskal-Wallis sum-rank test and unpaired Wilcoxon rank-sum test (LEfSE), with a p< 0.05.

## Results

### Host-microbiome interactions within the OTM: early response

We tested the hypothesis that oral tissue anatomical architecture together with salivary flow and composition (**Figure 1**) support long-term host-microbiome homeostatic interactions (three days and seven days post-inoculum; **Supplementary Figure 3** and **Figure 2**, respectively). We first established the importance of a native regime to preserve a healthy microbiome profile while promoting long term host viability. We inoculated a pool of healthy subgingival plaque microbiome (**Supplementary Figures 3-4**; day 0 post-inoculum) within the periodontal pocket of the OTM and monitored both host and microbiome early responses. The host demonstrated viability and functionality after three days, where the native culture regime supported a balanced ratio between proliferative (Ki67^+^) and differentiated (DAPI^+^/Ki67^-^) epithelium, suggesting epithelium robustness to microbial challenge. Moreover, we visualized bacterial adhesion and distribution through CLSM and SEM by processing the OTM in three regions (upper, medium, and lower) according to oxygen tension parameters (**Supplementary Figure 6**)^6^. Syto9 staining confirmed microbiome integration into the model and formation of biofilm bridges forming structural networks^37,38^, essential for biofilm formation. Relative abundance analysis (**Supplementary Figure 3**) confirmed the ability of the OTM, together with the properties of the microenvironment, to preserve richness and diversity, although decreased and segregated from the initial inoculate (*h*-plaque 0). Interestingly, *Neisseria* and *Fusobacterium*, two pivotal commensals and colonizers in biofilm formation^39^, were preserved and periopathogens, such as the genus *Porphyromonas* (*h*-plaque 0: 2%; upper-lower region: 3%), were present in limited abundance, suggesting that stimulation of salivary flow favored commensal genera and reduced the abundance of periopathogens. Hence, we demonstrated the importance of a native culture regimen to maintain a healthy microbiome, while supporting host tissue functionality.

**Figure 2.**
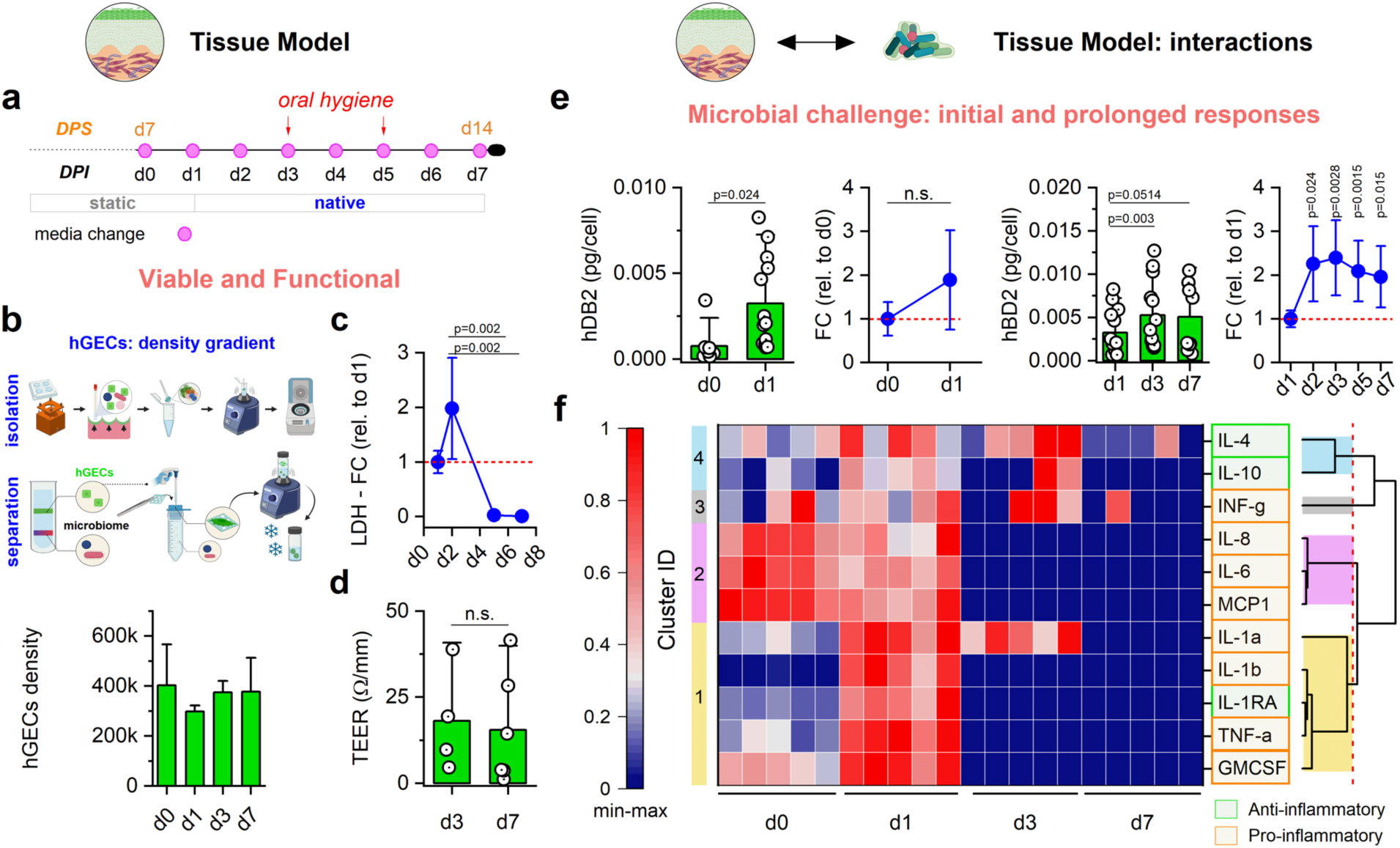
The tissue model remains viable, functional, and maintains homeostasis after microbial challenge. **a)** Mammalian-microbiome culture timeline: DPS: days post-seeding; DPI: days post-microbial inoculum. **b-d)** Viability and functional response of gingival tissue to microbial challenge. **b)** Cartoon created partially in biorender.com showing isolation of gingival epithelial cells from the anatomical scaffold using the density gradient centrifugation (Percoll) technique and related density quantification by picoGreen. The bar plot represents the mean±SD. **c)** Change in LDH from day 1 (red line) after microbial inoculation. Two-way ANOVA repetitive measures with Bonferroni’s post-hoc, p-values (<0.05) are shown in the figure (n=5). The line plot represents the mean ± SD. **d)** TEER analysis (Ω/mm) of oral epithelium challenged with subgingival microbiome and cultured under dynamic regimen for three to seven days (n=3/time point). Two-sample Student’s t test, p-value>0.05, n.s. = not significant. The bar graph represents mean±SD and single data points are shown. **(e-f)** Microbial challenge: initial and prolonged responses – **e)** Top, left to right: initial response (d0-d1), hBD2: normalization of ELISA data (pg) to the number of cells obtained by picoGreen analysis (n=13), Paired Samples t-test, p-value<0.05 is shown in the figure; fold change (FC) relative to day 0 (red line) after microbial inoculation (n=13), Paired Samples t-test, p-value>0.05 is shown in the figure, n.s. = not significant. Prolonged response (d1-d7), hBD2: normalization of ELISA data (pg) to the number of cells obtained by picoGreen analysis (n=13), One-way ANOVA repeated measures with a Dunnet’s post-hoc (relative to d1), p-values (<0.05) are shown in the figure (n=13). **f)** Heat-map showing host’s pro and anti-inflammatory cytokine levels over time, as indicated. Each cytokine has been rescaled with a min-max normalization. The dotted red line indicated the cut-off value. Each cluster is color-coded, as indicated (n=5).

### Host-microbiome interactions within the gingival tissue model: long-term response

Given the early eubiotic response host-microbe co-culture in combination with a sustained epithelial barrier, we investigated longer-term host-microbe interactions (**Figure 2a**). Epithelium stratification, assessed by density gradient coupled centrifugation (Percoll) and picoGreen techniques, showed sustained viability throughout the culture period, suggesting preservation of the multilayer (**Figure 2b** and **Supplementary Figure 5b)**. LDH analysis (**Figure 2c** and **Supplementary Figure 5c**) indicated an initial increase followed by a steady decrease in cellular cytotoxicity in the oral mucosa after inoculation. Barrier integrity (**Figure 2d**) was preserved up to seven days and no statistical changes were noted from the previous assessments at day three. Responses to microbial challenge were also investigated through the secretion of anti-microbial peptide and pro/anti-inflammatory factors (**Figure 2e-f**). Release of hBD2 normalized to cell number was found to increase with microbiome inoculum over time. Lastly, we monitored overtime the profile of cytokine release within the periodontal pocket upon inoculation (**Figure 2f**). After normalization against cell number (pg/cell), we performed a min-max normalization followed by a hierarchical clustering analysis on the cytokine’s temporal trajectory. We identified four clusters of cytokines with distinct temporal profiles, as reported in **Figure 2f**. In summary, cytokines within cluster 1, 3 and 4 were exhibited at lower levels before microbial challenge and displayed either transient increases (cluster1, except for IL-1α) or sustained elevation (cluster 3-4). On the other hand, cytokines in cluster 2 were initially expressed at higher levels at day 0, transiently decreased at day 1, and subsequently lacking. Hence, we demonstrated the maintenance and response of the oral epithelium within the OTM up to seven days and we characterized its interaction with polymicrobial populations.

### Subgingival plaque microbiome: viability and spatial organization within the OTM

To monitor microbial viability throughout the culture and its distribution within OTM, we performed viability assays and SEM analysis (**Figure 3**). Flow cytometry analysis (**Figure 3a** and **Supplementary Figure 7b)** showed a stable viable microbial population of the subgingival plaque microbiome from the time of the inoculum (79.1%) onto day 3 (66.6%) and day 7 (77.8%) post-inoculum, suggesting that a physiological saliva flow stimulation and oral hygiene procedures did not negatively impact the viability of the microbial population. Furthermore, viability was assessed by CLSM (**Figure 3b**). The microbial population, pre-stained with Syto9 before inoculation and stained with PI on day 7, appeared viable and homogeneously distributed within the depth of the scaffold. SEM micrographs showed the evidence of epithelium and a mature biofilm in all the regions, while different bacterial populations, such as cocci and rods, were identified in the upper and lower regions, respectively (**Figure 3b**). Therefore, we demonstrated that a physiologically relevant stimulation coupled with artificial saliva exposure favored microbial viability, distribution, and diversity.

**Figure 3.**
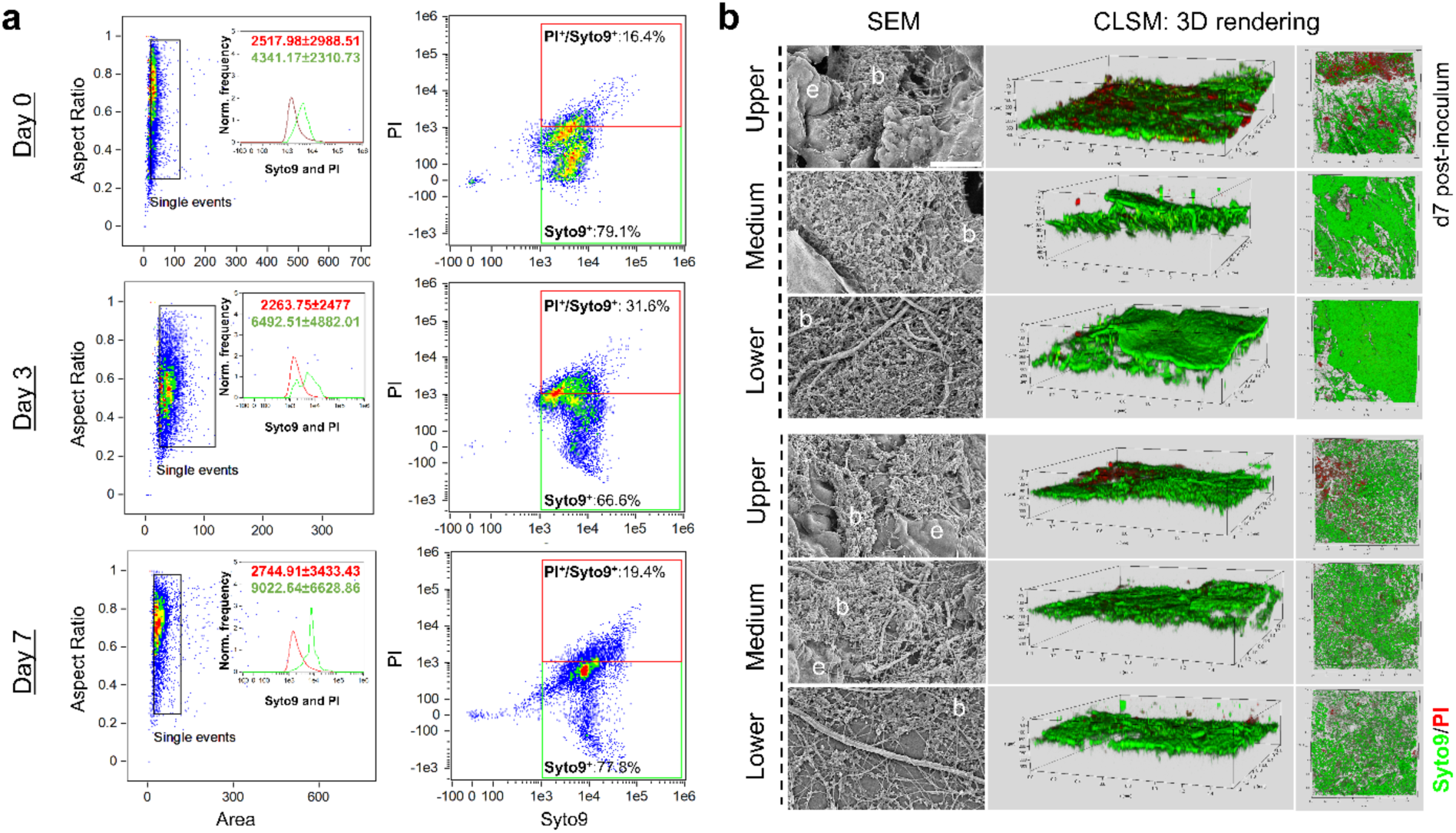
Sustaining a viable and complex human subgingival microbiome *in vitro*. Host-Microbiome interactions: **a)** Viability analysis of the microbial population before inoculation (day 0) and isolated from the anatomical scaffold by the density gradient centrifugation (Percoll) technique (day 3 and day 7). Left: RMS (Raw Mean Squared) values of the gradient and normalized frequency of the single event versus Syto9 (green) or Propidium Iodide (PI, red). The rectangle represents the individual events that were used for initial gating. Right: Flow cytometric analysis representing the percentage of positive microbial cells upon staining with Syto9 and PI. The percentages of cells expressing each dye are shown in the graph. **b)** SEM (n=2) and CLSM (n=5) analysis of microbial viability (Syto9/PI, green/red) and distribution within the entire depth of the scaffold. SEM scale bar = 10 µm. Labels: b: mature plaque biofilms; e = oral epithelium.

Tooth brushing and oral rising are considering effective daily procedures to minimize plaque buildup^40,41^. During host-microbiome co-culture within the OTM, oral hygiene was reproduced via oral rinse (mouthwash) for 10 seconds on day 3 and day 5 post-inoculum. We investigated potential negative effects of mouthwash exposure on the oral tissue viability and functionality, by monitoring host’s survival upon exposure to three commercial mouthwashes (Colgate®, Listerine®, and Crest®) at three different concentrations (**Supplementary Figure 1**). Live/dead assay (**Supplementary Figure 1a-b**) of Colgate® compared to the control (0%) showed statistically significant impairment on cellular viability starting from 2%. Although only Crest® at the highest concentration (8%) determined a statistically significant death compared to the control (0%), it showed a greater variability in the lower concentrations (1 to 5%). Listerine® maintained the highest percentage of cell viability under all the concentrations (p>0.05) for both gingival cells, and thus it was chosen as the mouthwash in our experiments. Lastly, given the effect of mouthwash on reducing plaque buildup and the nature of the scaffold in promoting cell adhesion, we performed a pilot experiment using Propidium Monoazide (PMA), as a pre-gDNA extraction method to ensure 16S rDNA sequencing of only living species at the time of the inoculum and after prolonged culture in the tissue model (**Figures 4-7 and Supplementary Figures 3,4,8-12**). To confirm gDNA amplification of only living microbial communities, subgingival plaque microbiomes were pooled, grouped in live and dead (heat-killed) samples, and treated with propidium monoazide (PMA) (**Supplementary Figure 2**). Electrophoresis results showed a band (500 bp) in both live samples PMA-treated (+) or not (-) and in dead samples not PMA-treated (-) with the photo-reactive DNA-binding dye. No band was detected in the dead PMA-treated (+) samples. These results confirmed the ability of PMA treatment, before gDNA amplification, in permanently modifying the DNA of dead bacterial cells, enabling genomic amplification of only live microbial cells, thus ensuring accuracy of the sequencing results.

**Figure 4.**
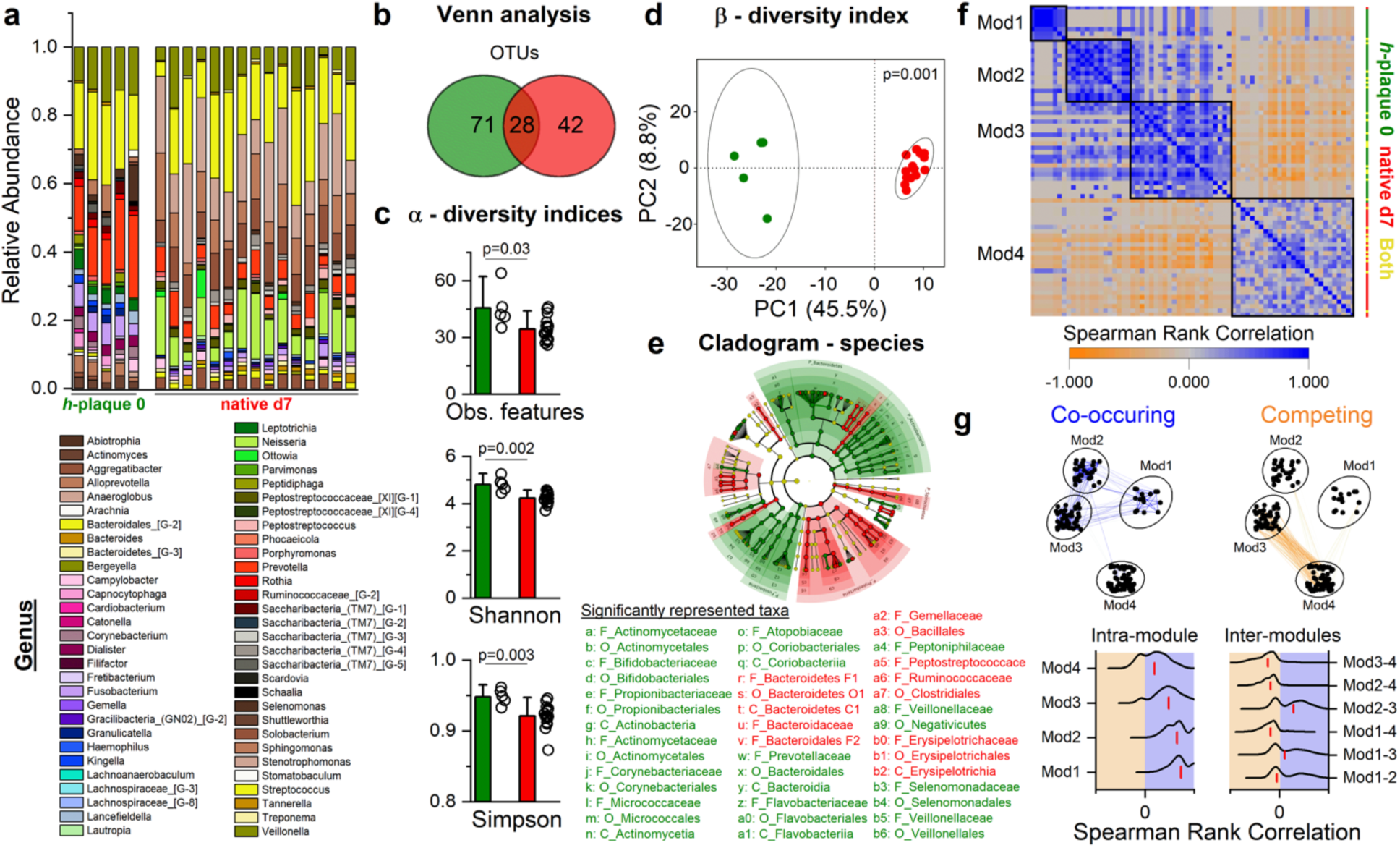
Physiological salivary flow promotes the growth of microbial communities and preserves most of their richness and diversity. Comparison between the original sample (green, N_patients_=8) and the microbiome cultured in native conditions (red, n=5) (*h*-plaque 0 *vs* native day 7) of: **a)** Relative Abundance – Genus; **b)** Venn analysis - species; **c)** Alpha-diversity indices: Observed features, Shannon and Simpson, Kruskal-Wallis H test, p-values (<0.05) are shown in the figure; **d)** Beta-diversity index plotted using Principal Component Analysis (PCA) by means of Euclidean distance (Aitchison distance), PERMANOVA test, p-value (<0.05) is shown in the figure; **e)** Differential abundance analysis shows statistically significant taxonomic levels by means of a cladogram-species (red and green nodes indicate statistically significant species, while yellow is not significant; the diameter of each circle is proportional to the abundance of the taxon represented). Names of statistically significant taxa are reported. **f)** Bacterial species similarity heatmap. Each black square indicated a distinct module. The color-coded vertical line on the right represented, for each species, whether the species was exclusively present at *h*-plaque 0 (green), at native day 7 (red), or present in both conditions (yellow). **g)** Top: Co-occurrence and competing networks (left and right, respectively). Edges were selected as greater than 0.7 or lower than -0.6 for clarity. Each node (black dot) represented a species, and each edge (line) represented the pairwise Spearman rank correlation between the connected nodes. Bottom: Edge distributions of intra- and inter-modules (left and right, respectively). Red vertical lines indicated the median value.

### Maintenance of healthy polymicrobial communities within the OTM

To evaluate the ability of the gingival tissue model to support the maintenance of a healthy, rich, and diverse microbiome, pooled subgingival samples were inoculated within the pockets of the model and cultured for seven days (**Figure 4**). Microbial communities were isolated from the three regions of the OTMs (**Figure 1**) on day 7 post-inoculum, sequenced and compared against the initial inoculate (*h*-plaque 0) (**Figure 4**). Analysis of relative abundance (**Figure 4a**) confirmed the presence of ubiquitous genera^28,42^ in the oral cavity, including *Peptostreptococcus, Streptococcus, Veillonella, Neisseria, Campylobacter, Fusobacterium Capnocytophaga or Prevotella*, supporting the claim of a diverse microbial community after seven days in culture. Compared to the initial inoculates (*h*-plaque 0), of 71 OTUs 28 were shared, while 42 were unique (**Figure 4b**). Diversity indices suggested a decrease in the overall observed OTUs (**Figure 4c**) and segregation from the original sample (**Figure 4d**). Lasty, the cladogram identified enriched phylotypes from *Bacteroidetes*, *Actinobacteria* and *Fusobacteria* for the *h*-plaque0 (green), while *Spirochete* and *Proteobacteria* were significantly represented in the OTM on day 7 post-inoculum (red) (**Figure 4e**). We then evaluated which species were more likely to co-occur or compete as a proxy of microbial social structure. Using the OTUs from *h*-plaque 0 and native day 7 post-inoculum, we built a pairwise similarity matrix (**Figure 4f**). Within this matrix, each entry represented a pairwise Spearman rank correlation (see **Supplementary Material_Modularity and similarity matrix** for complete list of pairwise values). Positive^43^ values (blue) denoted co-occurring species while negative values (orange) suggested competitive species; the magnitude of the correlation (absolute value) represented the strength of the association. We then performed a modularity analysis identifying four modules (or communities or niches). Each module represents groups of species that have strong positive correlations within the same module and weaker or fewer positive correlations with species belonging to different modules. We found that the modularity value, an index of the strength of the division of a network into communities, was equal to 0.48, indicating that the microbial network has a good level of structure, in which well-defined communities can be identified. Among the four identified communities, 1 to 4 displayed varying levels of within-module correlation strength, with module 1 exhibiting the strongest correlations and module 4 the weakest (**Figure 4g**, intra-module correlation distributions). Additionally, species within module 4 generally displayed an anticorrelation pattern with species from other modules (**Figure 4g**, inter-module correlation distributions). Further examination (**Figure 4f**, color-coded vertical line) revealed that modules 1 and 2 predominantly consisted of species abundant in the initial inoculate (about 87% of species), module 3 contained species present in both *h*-plaque 0 and native day 7 (25% of the species) with the remaining (75%) being present at *h*-plaque 0, and module 4 primarily comprised species identified only at native day 7 (78% of the species). In addition, we performed LEfSE analysis and hierarchical clustering, which showed several taxa highly enriched only in the initial sample or only at day 7 post-inoculum, while some of which were maintained throughout the culture (**Supplementary Figures 8-9**). Examples are *Streptococcus* or *Veillonella*, whose relative abundances were preserved in both groups, suggesting the favorable conditions of the model in supporting the growth of colonizers, while other colonizers, as *Actinomyces*^44^, did not survive. Enriched taxa on day 7 were identified as *Saccharibacteria TM7*^45,46^ or *Neisseria*^47^, which are health and host-associated bacteria, although the role of *Saccharibacteria TM7* is not fully understood yet.

Taken together, these data suggested that gingival anatomical model and mimicked stimulation of salivary flow promote the maintenance of subgingival plaque microbial communities and preserve most of the richness and diversity.

### Spatial organization of healthy polymicrobial communities within the OTM

Oxygen tension is a key environmental parameter within the periodontal pocket. Its gradient within the pocket provides the conditions for the co-existence of aerobic and anerobic species within the subgingival plaque^48,49^. To study the microbiome distribution throughout the pocket, we compared the microbial profile on day 7 post-inoculum of the upper, medium, and lower regions (**Figure 5**). Relative abundance (**Figure 5a**) of the genus *Streptococcus*, a facultative anaerobe^50^, was found greater in the middle (21%) and lower region (24,7%) than in the upper region (12%); *Neisseria*, an aerobic genus^51^, was in higher abundance in the upper region (14%), than in the middle (10%) and lower (7%) regions; *Veilonella*, an anaerobe^52^, showed a regional distribution decreasing from lower to upper (11 to 5%), respectively. About half of the species (54%) were distributed across all three regions, indicating bacterial colonization across the entire depth of the pocket. Several species were only detected in the upper region (15%), with the upper and medium region uniquely sharing 13% of the species and the upper and lower region less than 3%, thus suggesting a spatial distribution for a subset of the species analyzed, while maintaining overall richness and diversity (**Figure 5b-c**). Spatial distribution was also noticeable from β-diversity analysis (**Figure 5d**), from which the upper region segregated differently from both medium and lower, suggesting that the microbial profiles of the middle and lower regions (facultative or strictly anaerobes) belonged to the same cluster. Cladogram (**Figure 5e**) through differential abundance analysis identified enriched phylotypes from *Firmicutes and Saccharibacteria TM7* for the lower region (red) and *Proteobacteria* in the upper region (blue), while LEfSE (**Figure 5f**) showed 21 statistically significant taxa (LDA score >2) for the upper region, and 2 and 19 taxa for the medium and lower, respectively. Examples are *Gemella*, a commensal and facultative anaerobe genus^53^, present in higher percentage in the middle and lower regions, or *Aggregatibacter*, another facultative anaerobe genus^54^, highly enriched in the upper region, but also abundant in the other regions. Overall, analysis of anatomical gingival scaffold depth supported the distribution of the subgingival plaque microbiome, indicating regional differences with respect to oxygen tension.

**Figure 5.**
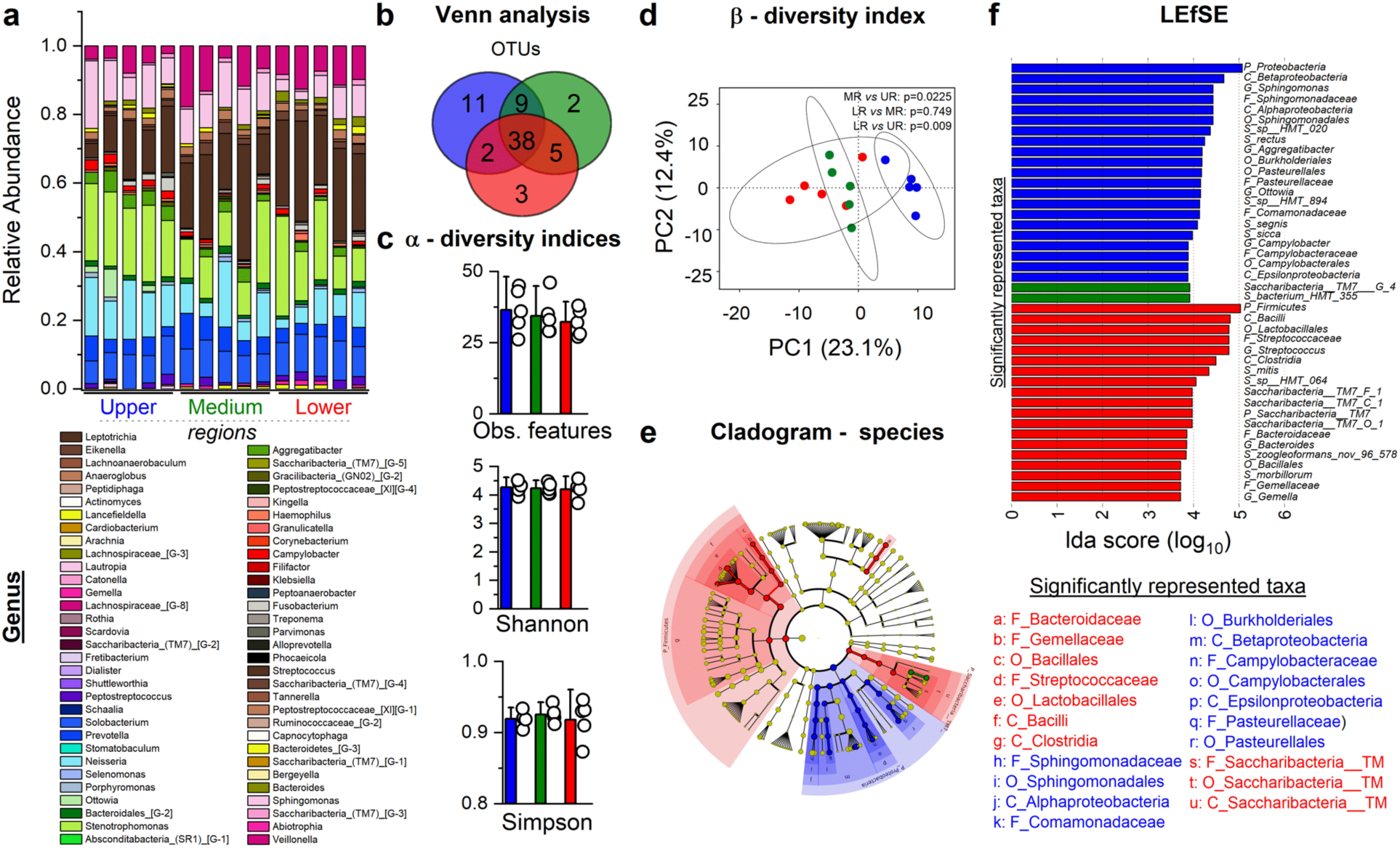
Construct anatomical architecture supports the organization of the human subgingival microbiome, indicating regional differences with respect to oxygen tension. Comparison among upper (blue), medium (green) and lower (red) regions of microbiome (n=5) cultured in the gingival bioreactor for seven days based on: **a)** Relative Abundance – Genus; **b)** Venn analysis - species; **c)** Alpha-diversity indices: Observed features, Shannon and Simpson; **d)** Beta-diversity index plotted using Principal Component Analysis (PCA) by means of Euclidean distance (Aitchison distance), PERMANOVA test, p-values (<0.05) are shown in the figure; **e-f)** Differential abundance analysis showing statistically significant taxonomic levels by means of a cladogram-species (red and blue nodes indicate statistically significant species, while yellow is not significant; the diameter of each circle is proportional to the abundance of the taxon represented) and taxa associated with statistically significant enriched bacteria between two groups by linear discriminant analysis Effect Size (LEfSE). Names of statistically significant taxa are reported in the figure.

### Intrinsic inter-variability of different healthy polymicrobial communities

The proposed model aims to incorporate the complex nature of microbial communities in the oral niche while maintaining the overall composition and dynamics of the microbiome, and thus with the host. However, microbial richness and diversity vary among individuals^55^, and incorporating such fingerprints *in vitro* requires different starting microbiomes in each experiment. To confirm inter-variability in the microbiome before inoculation in the anatomical model, patients-derived microbial communities (N_patients/pooling_=8) were pooled together, processed for 16S rDNA sequencing, and compared (**Figure 6**). Relative abundance (genus) between different initial pooled plaques (red and green) (**Figure 6a**) were also compared against the Human Oral microbiome Database^29^. This comparative analysis confirmed the presence of the ten most abundant genera in the subgingival plaque (ranked from the most to the least abundant): *Streptococcus, Fusobacterium, Prevotella, Capnocytophaga, Actinomyces, Corynebacterium, Rothia, Neisseria, Veillonella and Leptotrichia*. Venn analysis (**Figure 6b**) identified 69 shared OTUs, while 34 and 30 were specifics to each pooled group, respectively. The α and β-diversity indices (**Figure 6c-d**) indicated significant differences between the two original samples, supporting inter-variability in the microbial composition among individuals. Lastly, LEfSE analysis (**Figure 6e-f**) was performed to compare differential abundance between the two microbiomes. LEfSE revealed 80 differential abundant taxa with an LDA score higher than 2. A comparison (**Figure 6e-f**) between the two groups found 23 taxa enriched in the *h*-plaque 0 (green) group sample, whereas 57 were found in the other (*h*-plaque 0, red). In conclusion, although working with isolated subgingival plaques from human donors is crucial to replicate *in vivo* features, their high degree of variability should be recognized, emphasized, and used as a comparator in *in vitro* longitudinal studies.

**Figure 6.**
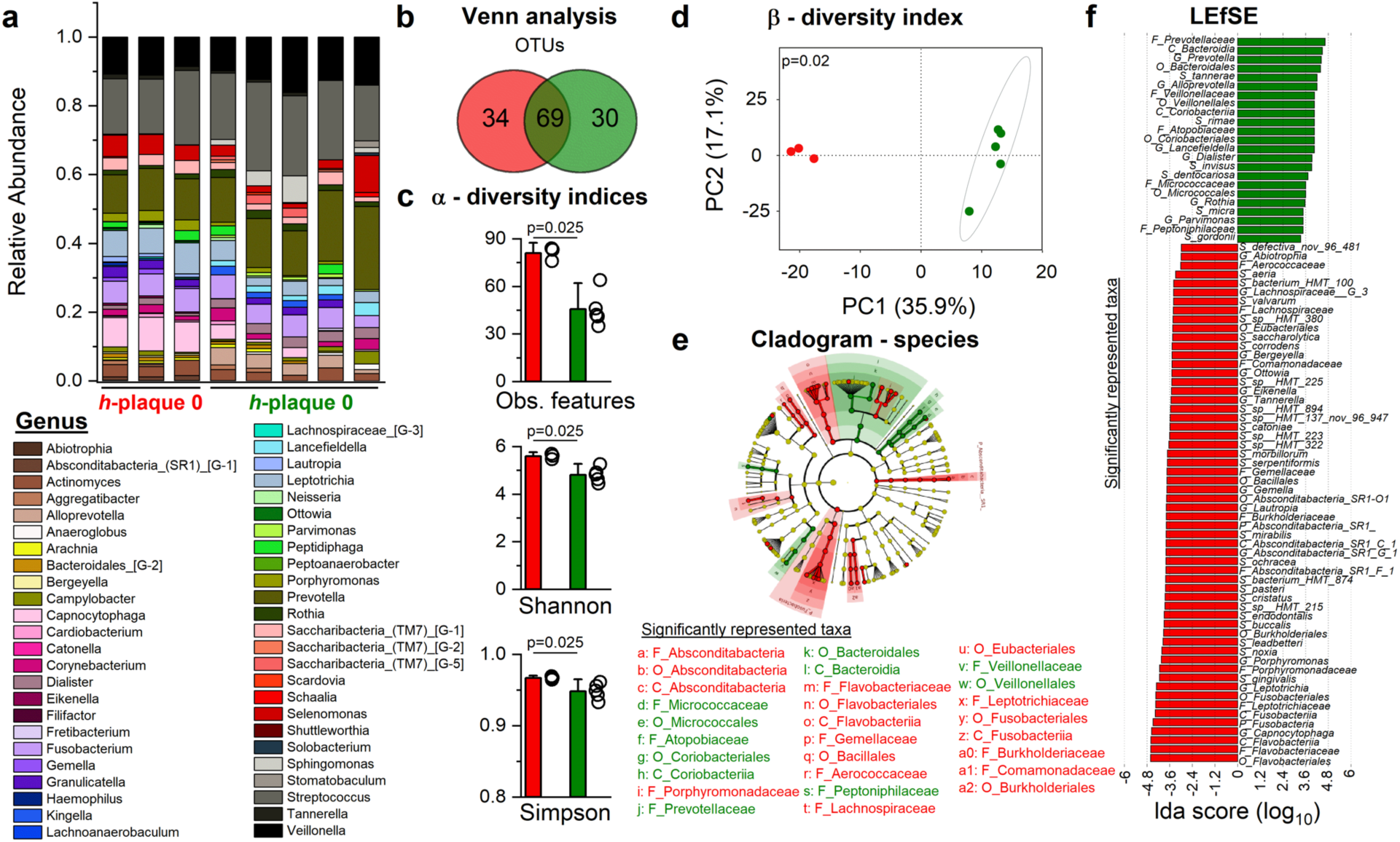
Intervariability of the human subgingival microbiome. Comparison between two isolations from human donors showed in red and green (N_patients_=3-8) of: **a)** Relative Abundance – Genus; **b)** Venn analysis - species; **c)** Alpha-diversity indices: Observed features, Shannon and Simpson, Kruskal-Wallis H test, p-values (<0.05) are shown in the figure; **d)** Beta-diversity index plotted using Principal Component Analysis (PCA) by means of Euclidean distance (Aitchison distance), PERMANOVA test, p-value (<0.05) is shown in the figure; **e-f)** Differential abundance analysis showing statistically significant taxonomic levels by means of a cladogram - species (red and green nodes indicate statistically significant species, while yellow is not significant; the diameter of each circle is proportional to the abundance of the taxon represented) and taxa associated with statistically significant enriched bacteria between two groups by linear discriminant analysis Effect Size (LEfSE). Names of statistically significant taxa are reported in the figure.

### Eubiosis within the gingival tissue model

To validate the maintenance of an eubiotic condition within the periodontal pocket *in vitro*, we analyzed α-diversity indices over-time (**Figure 7**). Compared against initial inoculate, each region at day 3 post-inoculum showed a statistical difference for each index (**Figure 7a**), while similar richness and uniformity were identified after seven days in culture (p>0.05). Analysis of clustered regions *versus h*-plaque 0 (**Figure 7b**) showed a statistically significant decrease in all three α-diversity indices for the day 3 post-inoculum OTMs, while only the observed features were preserved (p>0.05) for day 7 post-inoculum OTMs. Notably, despite the differences in the static condition, the α-diversity indices increased from day 3 to day 7 post-inoculum, strongly suggesting the ability of the system to recover from the initial decrease in diversity and thus corroborating the maintenance of a homeostatic and eubiotic state in the proposed model.

**Figure 7.**
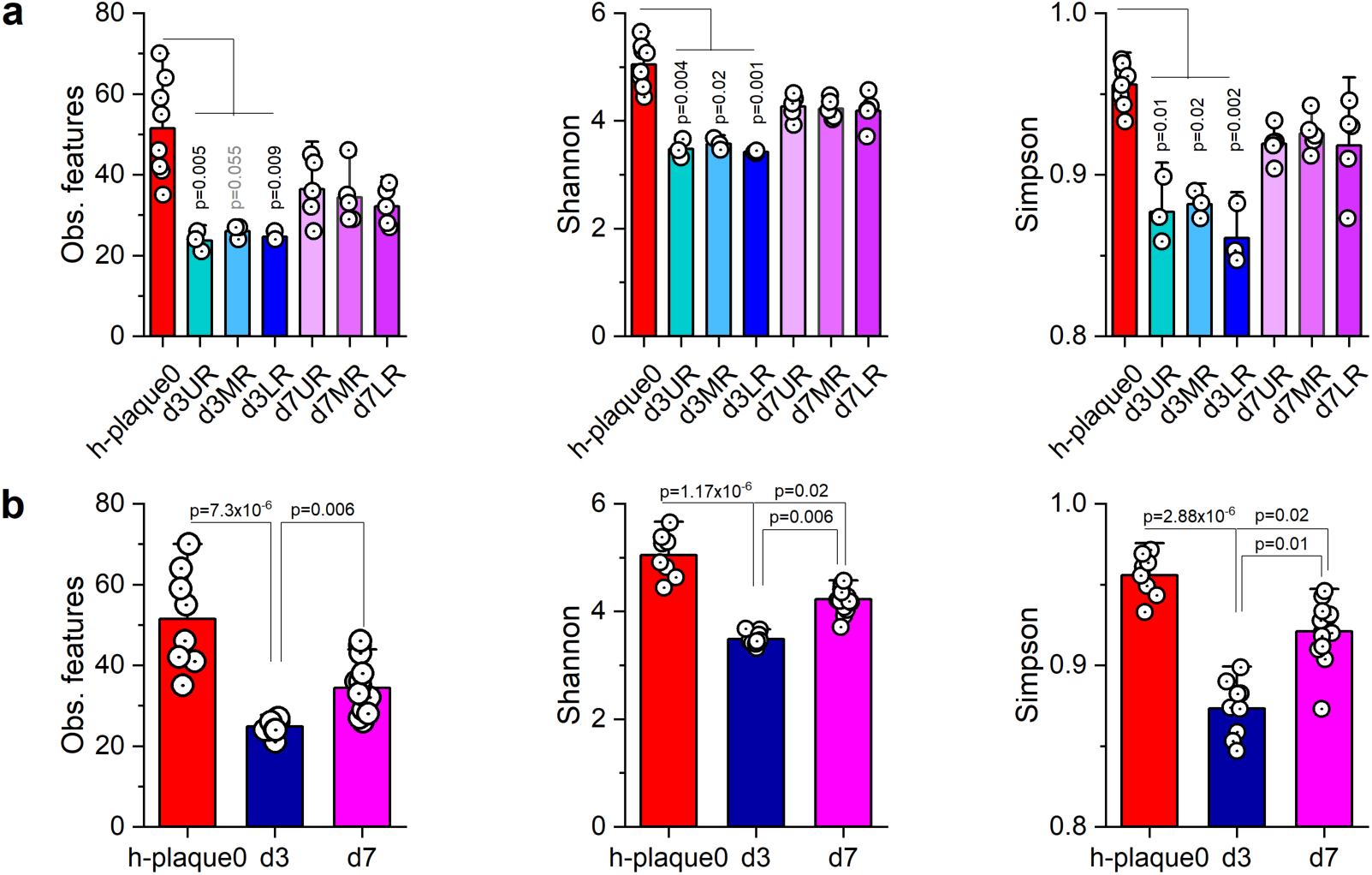
Prolonged culture equilibrates richness and diversity indices within the anatomical construct. **a)** Comparisons of alpha diversity indices between the original inoculate (*h*-plaque 0, red) and the upper (light blue, n=3), middle (blue, n=3) and lower (dark blue, n=3) regions cultured for 3 days or the upper (light pink, n=5), middle (pink, n=5) and lower (dark pink, n=5) regions cultured for 7 days; Kruskal-Wallis H test with Dunn’s test for non-parametric pairwise comparisons, p-values (<0.05) are shown in the figure. **b)** Comparisons of alpha diversity indices among the original sample (*h*-plaque 0, red), samples at day 3 (royal blue) and at day 7 (magenta); Kruskal-Wallis H test with Dunn’s test for non-parametric pairwise comparisons, p-values (<0.05) are shown in the figure.

## Discussion

Long-term interactions between host and oral microbiome have yet to be fully replicated in an *in vitro* system. *In vitro* experimental solutions must provide the native cyto-anatomical architecture, saliva buffering, native fluid dynamic regime, as well as physiological oxygen/pH gradients that, collectively, we have shown to support a homeostatic relationship between host and microbiome for at least seven days. While we previously seeded human subgingival plaque microbiome in a three-dimensional replica of the human gingiva for 24h^6^, in the present manuscript we advanced our OTM and extended the host-microbiome co-culture to seven days by introducing a salivary native dynamic regime^10^. Given the complexity of microbial and host factors that contribute to periodontal health, it is necessary to fully comprehend how the host and microbial communities communicate, interact, and adapt within our *in vitro* model. We were able to investigate both the host and the microbiome with a range of techniques including, but not limited to, cytokine profiles, epithelial barrier function, cell viability, 16S rDNA sequencing and microbial sociability.

One of the main challenges in oral tissue engineering is to maintain host-microbiome cultures beyond 24h, due to metabolic competition between microbial and mammalian communities^2,56^. To support the maintenance of host-microbiome equilibrium, we also performed mouthwash rinses, as commercial oral rinses are known to reduce microbial load^40^. Commercially available mouthwashes, such as Crest®, Colgate® or Listerine®, contain active components such as Cetylpyridinium chloride (CPD), Chlorhexidine (CHX) gluconate- or essential oils (eucalyptol, menthol, methyl salicylate and thymol) to reduce bacterial viability^40^. To the best of our knowledge, the use of mouthwash in a physiologically relevant model with the purpose of preserving the host while reducing microbial load has never been attempted before. Our data (**Supplementary Figure 1**) indicated that Colgate® and Crest® impaired the viability of primary stromal cells, while Listerine® showed no significant effect on either gingival cell types. Accordingly, rinses with diluted Listerine (1% v/v) for 10 seconds were performed on day 3 and day 5 post-inoculum, which as shown by host ‘s analysis helped preserve cell viability and oral barrier (**Figure 2**). In our model, the host was shown to be still viable (**Figure 2b-c**) and functional (**Figure 2d**) after seven days of co-culture with the human oral microbiome. We were also able to assess the host DNA content (picoGreen), a proxy for cell quantification, without contamination by prokaryotic DNA. Indeed, by adapting a density gradient centrifugation method (Percoll®)^23^, we separated eukaryotic from procaryotic cells and analyzed the DNA content of the eukaryotic fraction. Furthermore, TEER measurements showed that the host barrier was not affected by co-culture with the microbiome, suggesting the integrity of the epithelial barrier in the face of healthy microbial challenge.

An important feature of the host is its ability to respond dynamically to the presence of the microbiome. *In vivo*, this is ensured by the release of secreted factors such as antimicrobial peptides and cytokines^57^. Human beta-defensin 2 (hBD2) is a crucial antimicrobial factor released by epithelial cells aimed at buffering bacterial abundance and fighting periopathogen overgrowth^58^. Release of hBD2 is also promoted by oral commensals such as *Fusobacterium nucleatum*^59,60^. In response to the bacterial challenge, our OTM showed persistent production of the antimicrobial factor hBD2 (**Figure 2e**), further sustaining an active host response to the microbiome targeted at preserving host-microbiome equilibrium^2,13^. The host also released anti- and pro-inflammatory cytokines with distinct temporal profiles (**Figure 2**). We identified (**Figure 2f**): 1) cluster 1 (pro-inflammatory: IL-1α, IL-1β, TNF-α, GMCSF; anti-inflammatory: IL-1RA) composed of cytokines expressed at low levels before the microbiome inoculum that increased transiently at day1 (with the exception of IL-1α); 2) cytokines in cluster 2 (pro-inflammatory: IL-8, IL-6-MCP1) were already released at day0 and, after a drop in their levels at day1, were subsequently not detected; 3) cluster3 (anti-inflammatory: INF-γ) and cluster4 (anti-inflammatory: IL-4, IL-10) were composed of cytokines that were present at day 0 and increased their levels after the microbial challenge for all seven days. Of note are the pro-inflammatory cytokines IL-1α-β, secreted by epithelial and stromal cells after being stimulated by microbial population, which are pivotal in the recruitment of immune cells^57,61^ and whose levels are stabilized in tissue homeostasis by anti-inflammatory cytokine IL-1Ra^61^. Accordingly, we found these cytokines clustered together (cluster1). In addition, it is important to notice that IL-1α-β are only transiently upregulated at day1; in fact, excessive and prolonged secretion of IL-1α-β causes collagen degradation and bone resorption in the periodontium^61^. In addition, IL-4, an anti-inflammatory cytokine, was detected throughout the culture period. It has been reported that IL-4, by supporting anti-inflammatory action, is able to downregulate pro-inflammatory cytokines^57^, which could explain the non-detection after day 1 of pro-inflammatory cytokines and the overall restoration of homeostasis. Hence, these data suggest that our system could be integrated with immune cells in the future, given the host’s ability to actively signal microbial presence and mimic physiological responses through pro- and anti-inflammatory cytokines.

In our model, we employed human subgingival plaque microbiome rather than the co-culture of selected microbial species. In fact, microbial species richness is instrumental in establishing a physiological environment characterized by host-microbiome and microbe-microbe interactions. Furthermore, one of the key findings of the polymicrobial synergy and dysbiosis ( PSD) model is that a single keystone pathogen is unable to initiate dysbiosis unless aided by other microbial species within the community^62,63^, further supporting the need of diverse and complex social interactions to properly mimic these processes. Before proceeding to sequencing analysis, we assessed microbial viability before inoculum (initial inoculate/h-plaque 0) and in the *in vitro* OTM. The microbiome was not only viable after harvesting (**Figure 3a**- 79.1% viable cells) but remained viable during the seven days of culture (day7: 77.8% viable cells). The factors that mostly contributed to the preservation of viable microbial cells were the establishment of local oxygen gradient at the periodontal niche and the pH buffering capacity of the saliva, which together ensured the stability of a physiological pH and the preservation of a healthy and eubiotic microbiome^2,10^ (**Supplementary Figure 6**). CLSM and SEM further showed the development of thick layers of microbial population within the entire depth of the anatomical scaffold, which may suggest mature biofilms^17,46^. Biofilm initiation can be more clearly appreciated in day 3 post-inoculum OTMs samples, while the multi-layer organization of bacterial cells at day 7 post-inoculum makes visualization of the biofilm base more difficult^17^.

We then evaluated the microbial species present in the human plaque (initial inoculate/*h*-plaque 0) and after three or seven days (native day 3 or native day 7 post-inoculum) of co-culture in our OTM. To eliminate DNA contamination of dead cells during sequencing, we pre-treated the analyzed microbiome (*h*-plaque 0, native day 3, native day 7 post-inoculum) with propidium monoazide (PMA) (**Supplementary Figure 2**). PMA is a DNA-intercalating dye, which binds the small groove of the DNA in dead cells, and, upon blue LED light photo-activation, modifies the DNA so that it cannot be amplified in PCR^64–66^. The OTM on day 7 post-inoculum sustained microbial diversity and confirmed the abundance of healthy-associated and commensal genera (**Figure 4**) such as *Streptococcus*, *Neisseria*, *Veillonella*, *Gemella, Leptotrichia,* some of which highly enriched on day 7 compared with *h*-plaque 0 (**Supplementary Figures 8-9**). Although a decrease in α-diversity indices compared with *h*-plaque 0 was detected, we observed an increasing trend from day 3 post-inoculum (**Figure 7**) in richness and diversity, supported by Cohen’s effect size (*d*) (**Supplementary Table 1**: *h*-plaque 0-native day 7 (*d*_obs.feat_:1.24; *d*_Shannon_: 2.13; *d*_Simpson_:1.88) compared with hplaque0-day 3 (*d*_obs.feat_:6.46; *d*_Shannon_: 12.3; *d*_Simpson_:7.53). The decreased effect size of the *h*-plaque 0-native day 7 post-inoculum comparison compared with *h*-plaque 0-native day 3 post-inoculum suggested that the OTM was able to recover from the initial loss in diversity because of longer culture periods, as reported in other *in vitro* investigations^67,68^. Consistent with this result, on day 7 post-inoculum (**Figure 5**), but not on day 3 post-inoculum (**Supplementary Figure 4**), there was a spatial distribution of species within the depth of the pocket; in particular, the upper region was found to be different from the medium and lower regions, suggesting species segregation based on upper-related features (*i.e.,* oxygen, salivary shear), further supporting the need for long-term culture to replicate physiological characteristics and replicate spatial distribution *in vitro*. Moreover, prolonged culture periods favored the culture of genera, such as *Gemella* or *Peptostreptococcaceae[XI],* oral commensal species^69,70^, which were below the detection threshold (rel. abundance %) on *h*-plaque 0, indicating the ability of the model to promote the growth of low-abundance commensals at similar physiological abundance percentage and thus restore diversity^28^.

Importantly, the model preserved microbial social interactions. We identified four modules representing distinct bacterial communities (**Figure 4**), which supported the establishment of interbacterial dialogues within the OTM and further substantiating the need to utilize the whole microbiome to replicate social networking in the periodontal niche. For example, *Veillonella parvula*, which is crucial in the colonization of the niche as a bridging taxon in sustaining the growth of multispecies in the biofilm, was found to be highly correlated with the species *Streptococcus mitis* (Spearman’s rank correlation value of 0.66) (**Figure 4**), suggesting not only biofilm architecture, but also the regulation of community metabolisms and thus the preservation of a health ecosystem^71–73^. Lastly, although microbial communities and functions are shared, the composition of the microbiome is largely and highly individualized based on environmental and host factors^9,55,63^. Despite an evident inter-variability (**Figure 6**) among samples collected at *h*-plaque 0, our model allowed us to show robust and consistent effects, which align with physiological conditions at the microbiome-host interface.

In conclusion, we advanced our three-dimensional humanized gingiva model^6^ by co-culturing it with healthy human subgingival plaque microbiome for seven days within an oral bioreactor that mimics native salivary dynamics^10^. Our results indicated sustained viability of host and microbial communities, native topographical phenotypes and physiological response, preservation of healthy microbial populations and interbacterial dialogues. The model can be also useful in the identification of biomarkers associated with eubiotic/dysbiotic profiles benchmarked against clinical investigations. Future studies will be based on further advances in the model, as it lacks periodontium components (*i.e.,* alveolar bone, neutrophils and macrophages or vasculature), which are instrumental in understanding the trajectories of periodontal inflammation.

## Acknowledgements

We would like to thank the National Institute of Dental & Craniofacial Research of the National Institutes of Health (R03DE030224) and ORAU (Ralph E. Powe Junior Faculty Enhancement Awards) for supporting this work. We are thankful to all the students who have contributed to this project - Devon R. Hartigan, Eric R. Webb and Helen B. Hurley - and to Tsute Chen, PhD, for his help in navigating the sequencing reports. Additionally, we would like to thank Mattia Bonzanni, PhD and B. J. Black, PhD for their input on the manuscript.

## Author contributions

M.A.: Methodology, validation, formal analysis, investigation, data curation, visualization, writing – original draft preparation, writing – review and editing. G.E.C., A.R.D., H.H., B.J.P.: Methodology, formal analysis, writing – review and editing. X.H.: Writing – review and editing. C.E.G.: Conceptualization, methodology, validation, data curation, writing – original draft preparation, writing – review and editing, supervision, project administration, funding acquisition. All authors have read and agreed to the published version of the manuscript.

## Data availability

The main data supporting the findings of this study are available in the Article and Supporting Information. The raw data generated in this study are available from the corresponding author on reasonable request.

