## Supplemental Figures for "Sustaining healthy long-term host-microbiome interactions in a physiologically relevant dynamic gingival tissue model"

#### Supplementary Figures

#### Supplementary Figure 1

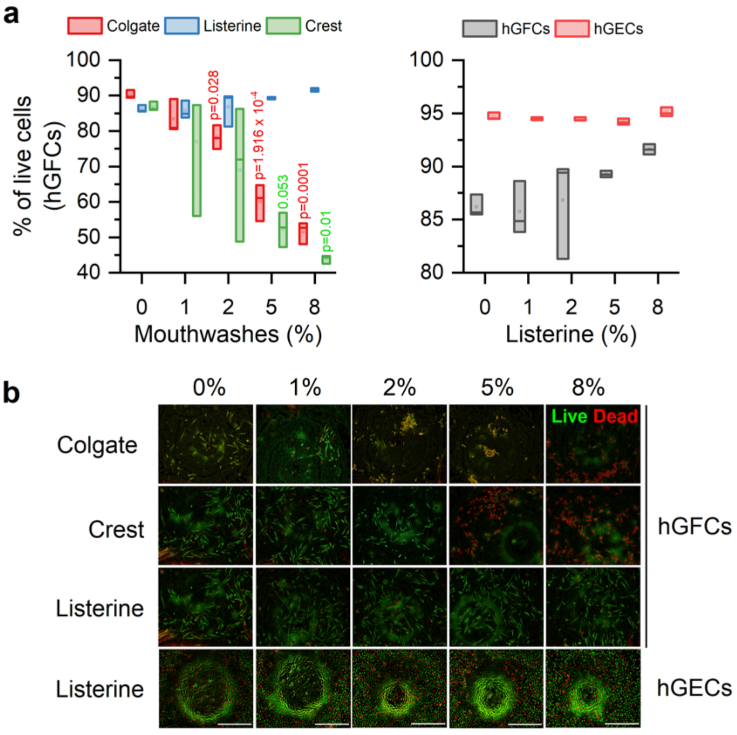


**Supplementary Figure 1. Effects of oral hygiene mimicking on the gingival tissue model. a)** Live/dead analysis of hGFCs exposed to different commercially available mouthwashes (left) and exposed to different concentrations of Listerine (right - 0, 1, 2, 5, 8 %). One-way ANOVA relative to 0% with Bonferroni's post-hoc, p-values (<0.05) are shown in the figure (n=3). **b)** Immunofluorescence composites (Live, green / Dead, red) of hGFCs (n=3/condition) and hGECs (n=3/condition) exposed to increasing concentrations of mouthwashes (in %: 0, 1, 2, 5, 8).

**Supplementary Figure 2**

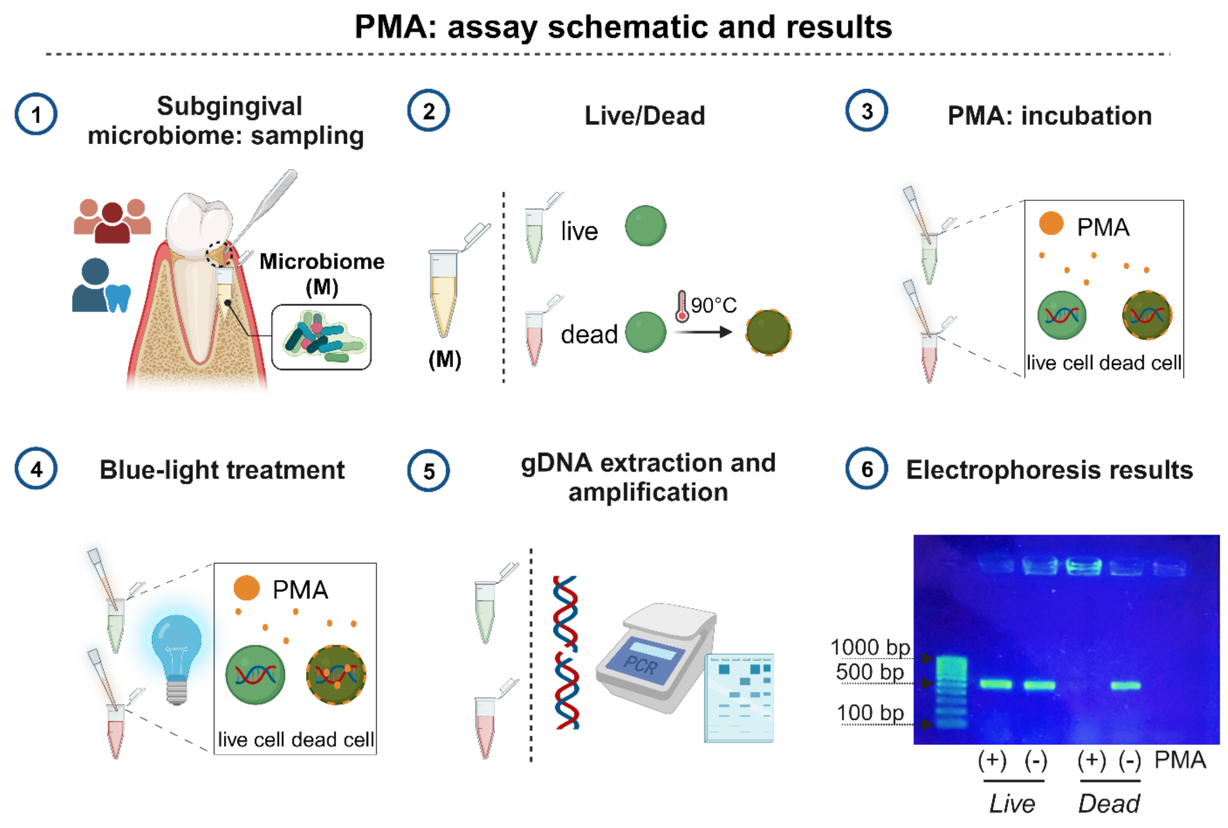


**Supplementary Figure 2. Propidium monoazide (PMA) treatment for accuracy of 16S rDNA sequencing analysis.** Schematic (steps:1-6), created in biorender.com, of the workflow of propidium monoazide (PMA) treatment for human subgingival microbiome. Step 6 shows electrophoresis gel of PCR endpoint products (16S rRNA gene: V1-V3 variable regions) pointing at the accuracy of PMA (band showed at 500 bp) in permanently modifying the DNA of the bacterial population ($N_{patients}$=4) to avoid DNA amplification from dead cells.

#### Supplementary Figure 3

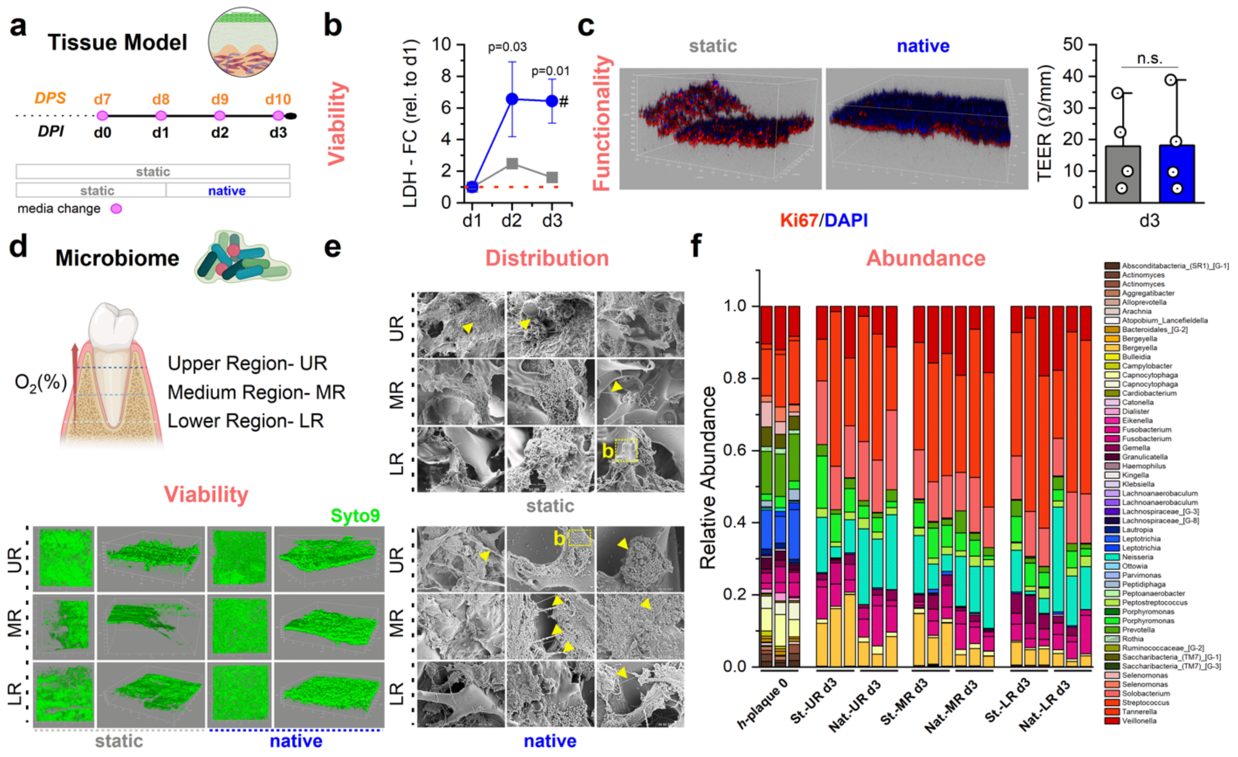


**Supplementary Figure 3. Native culture conditions to support a complex human subgingival microbiome without adversely affecting the host: early-response. a)** Mammalian-microbiome culture timeline. DPS: days post-seeding; DPI: days post-microbial inoculum. **b)** Host's viability analysis: change in LDH from day 1 (red line) after microbial inoculation between static (gray) and native (blue) conditions. Two-way ANOVA repeated measures with Bonferroni's post-hoc, p-values (<0.05) of the statistical differences between the two conditions are shown in the figure (n=3/conditions); hashtag (#) indicates the statistical difference (p-value) within the same condition overtime: native: d1 *vs* d2: 0.006; d1 *vs* d3: 0.007. Data shown in line plot represents mean±SD. **c)** Host's functional analysis: left, CLSM of stratified and differentiated epithelium identified by DAPI (differentiated, blue) and Ki67 (proliferative, red) staining on day 3 (n=2); right, TEER analysis (Ω/mm) between static (gray) and native (blue) conditions (n=3/conditions). Two-sample Student's t-test, p-value>0.05, n.s.= not significant. Data shown in the bar graph represents mean±SD, while circles represent single data points. Microbiome assessments (**d-f**): **d)** Viability: top, gradient of plaque organization-left, cartoon, created in biorender.com, showing gingival scaffold processing in three regions (upper, medium, and lower), previously identified by oxygen measurements (**Supplementary Figure 6**); bottom, CLSM (n=2/condition) analysis of microbial viability (Syto9, green) and distribution within the entire depth of the scaffold. **e)** Distribution: SEM (n=2/condition) analysis of microbial distribution within the entire depth of the scaffold. Scale bars (SEM) shown in the figure: 5 and 10 µm. Labels: yellow arrows indicate mature plaque biofilms and connecting biofilms; b = initialization of biofilms. **f)** Relative abundance – genus: comparison between the original sample (hplaque0) and gingival tissue model regions (upper, medium, and lower) cultured under static (n=3) or native (n=3) conditions and isolated after 3 days post-inoculum.

**Supplementary Figure 4**

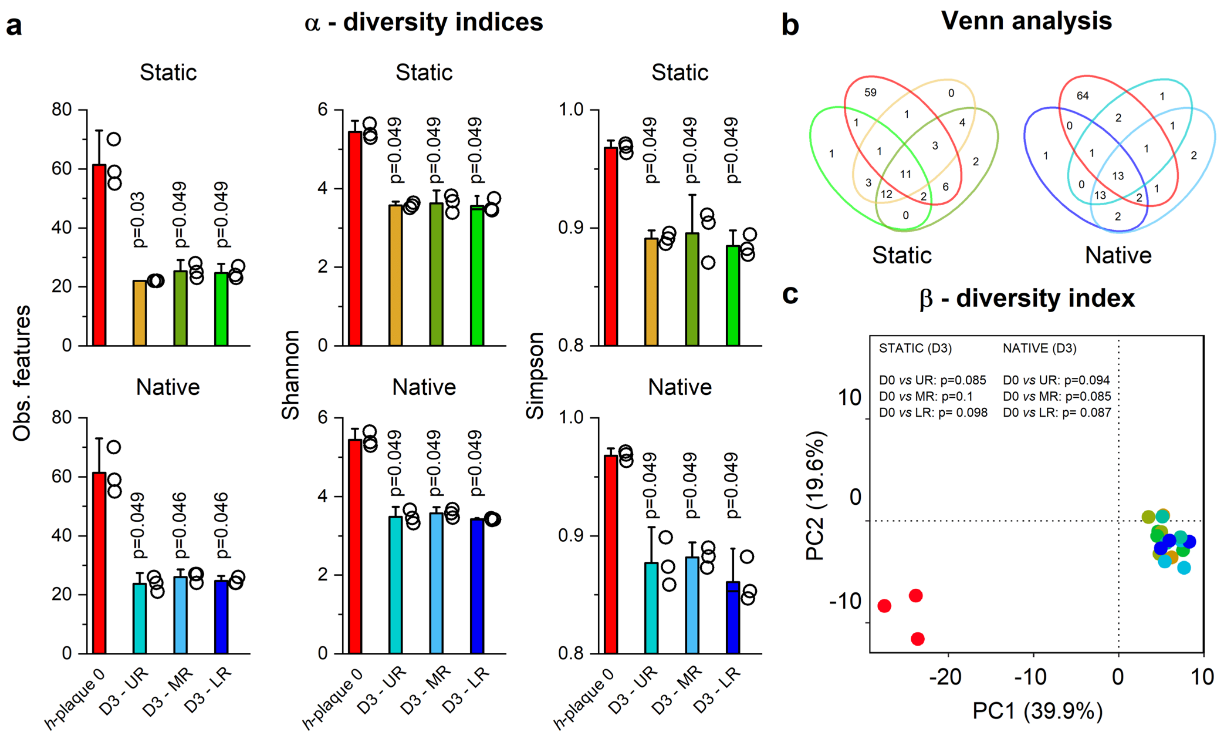


**Supplementary Figure 4. Analysis of the 16S rDNA alpha diversity indices of human subgingival microbiome cultured in artificial saliva in static *vs* native regimen: early-response.** Comparison between the original sample (*h*-plaque 0, red) and the upper (yellow, n=3), middle (dark green, n=3) and lower (green, n=3) regions cultured for 3 days under static regime or the upper (light blue, n=3), middle (blue, n=3) and lower (dark blue, n=3) regions cultured under native regime for 3 days (16h static + 2 days native) from the gingival tissue model based on: **a)** Alpha-diversity indices: Observed features, Shannon and Simpson, Kruskal-Wallis H test relative to hplaque0, p-values ($<0.05$) are shown in the figure; **b)** Venn analysis - species; **c)** Beta-diversity index plotted using Principal Component Analysis (PCA) by means of Euclidean distance (Aitchison distance), PERMANOVA test, p-values ($<0.05$) are shown in the figure.

**Supplementary Figure 5**

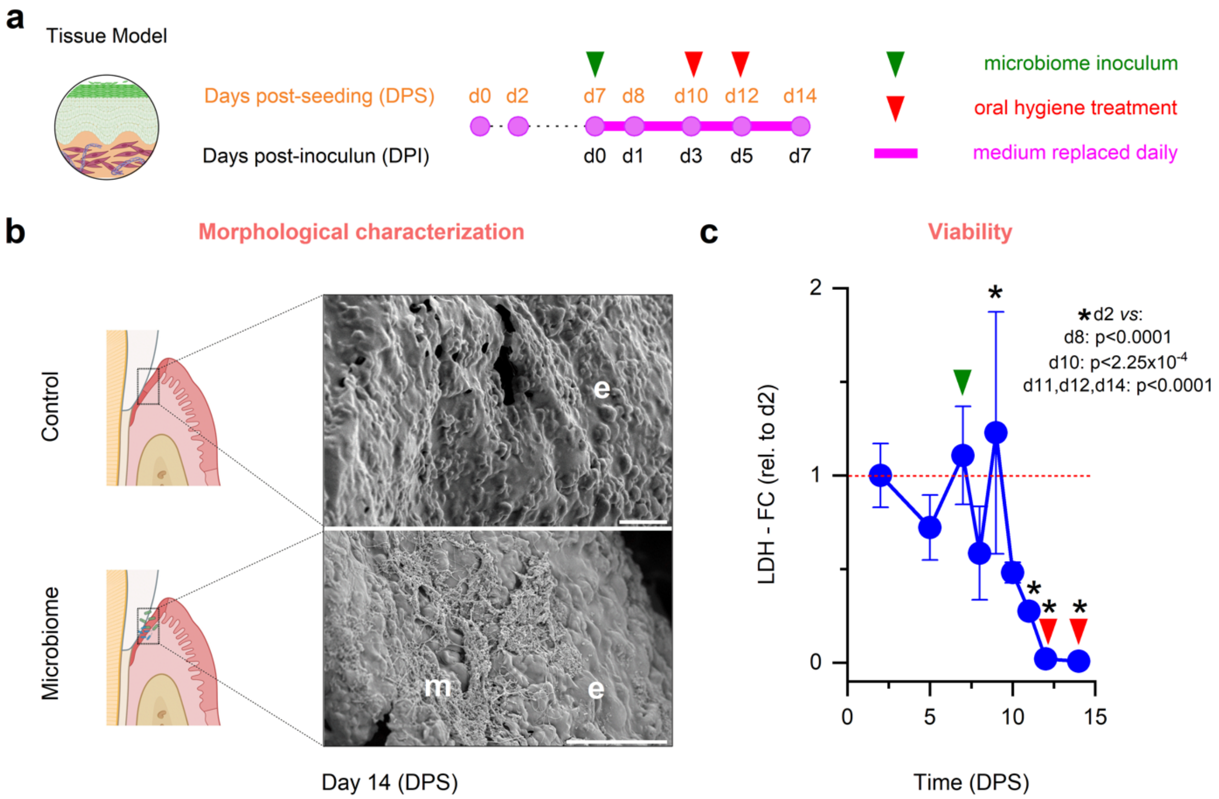


**Supplementary Figure 5. Effect of human subgingival microbiome on gingival tissue viability. a)** Host-microbiome culture timeline. **b)** SEM micrographs (n=2/condition) showing the gingival epithelium (**e**) in absence or in presence of microbiome (**b**) on day 14. Scale bar (upper to bottom): 20 and 30µm. The cartoon was created on biorender.com. **c)** Change in LDH (red line) relative to day 2 post-seeding. One-Way ANOVA repeated measures with Dunnett's post-hoc, p-values (<0.05) are shown in the figure (n=5). The line plot represents mean±SD.

**Supplementary Figure 6**

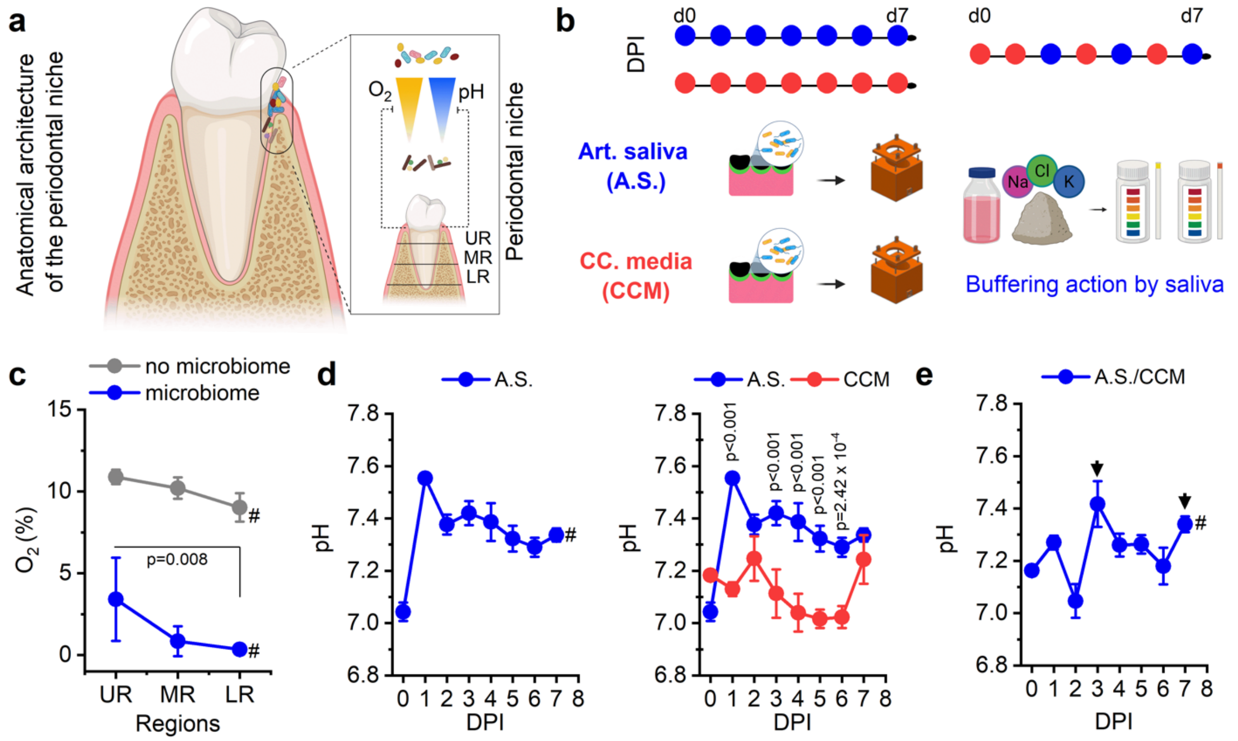


**Supplementary Figure 6. Artificial saliva oxygen tension and buffering action within the gingival tissue model.** a) Schematic of the anatomical architecture of the periodontal niche with emphasis on oxygen tension and pH readings. **b)** Mammalian-microbiome culture timeline based on different tissue culture media. **c)** Oxygen tension measurements ($O_2$%) in presence (blue) or absence (grey) of oral microbiome according to upper (UR), medium (MR), and lower (LR) regions in anatomical scaffolds. Two-Way ANOVA repeated measures with Bonferroni's post-hoc, p-values (<0.05). # indicates the significant statistical difference (no microbiome *versus* microbiome) within the same region: p-values<0.0001 for each comparison. The line plot shows mean±SD. **d)** Left, pH trajectory of the gingival tissue model cultured in artificial saliva. # One-way ANOVA repeated measures with Dunnet's post-hoc (relative to d1); p<0.05; p-values are reported in **Supplementary Table 2**. Right, pH trajectory of the gingival tissue model cultured in artificial saliva (blue) or co-culture media (red). Two-way ANOVA repeated measures with Bonferroni's post-hoc; p<0.05 are reported within the graph. The line plots show mean±SD. **e)** pH trajectory of the gingival tissue model cultured in alternating co-culture media and artificial saliva. # One-way ANOVA repeated measures with Bonferroni's post-hoc; p<0.05; p-values are reported in **Supplementary Table 3**. The line plot shows mean±SD. Arrows indicated the buffering actions of artificial saliva. Cartoons were partially created in biorender.com.

#### Supplementary Figure 7

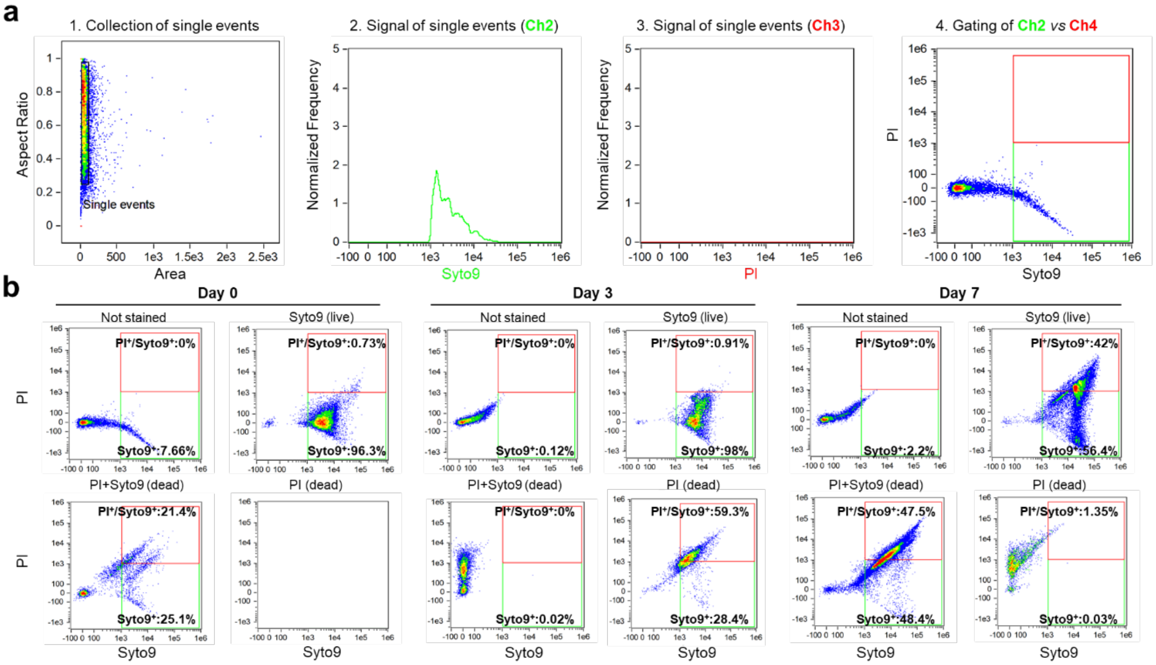


**Supplementary Figure 7. Flow cytometry analysis: parameters optimization. a)** Workflow can be summarized into four steps: 1) Collection of single events; 2-3) Fluorescent signal of single events corresponding to each dye; 4) Analysis (gating) of signal of Syto9 (live) *versus* Propidium Iodide (PI, dead). **b)** Flow cytometric analysis of controls representing the percentage of microbial cells positive for staining with Syto9 or PI before inoculation (day 0) and isolated from the anatomical scaffold by the density gradient centrifugation (Percoll) technique (day 3 and day 7). The percentages are shown in the graph.

**Supplementary Figure 8**

Significantly represented taxa

**hplaque0** *vs* native d7

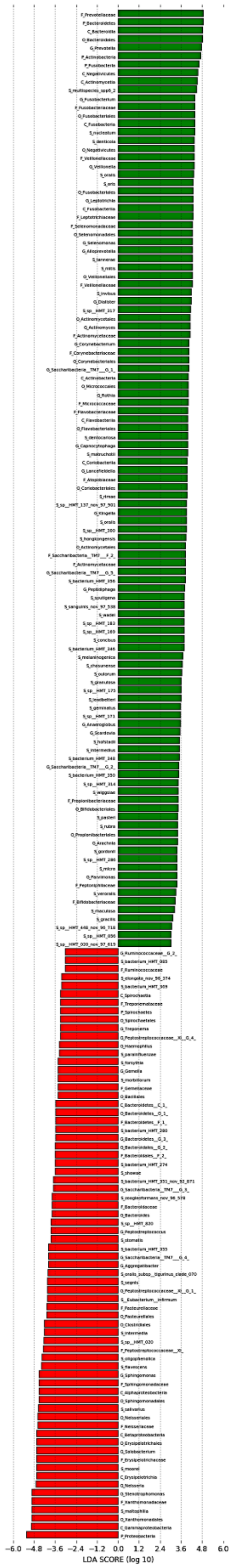

**Supplementary Figure 8. Differential abundance analysis (LEfSE) of human subgingival microbiome within the gingival tissue model: *h*-plaque 0 *versus* native d7.** Differential abundance analysis showing taxa associated with statistically significant enriched bacteria between two groups by linear discriminant analysis Effect Size (LEfSE). Names of statistically significant taxa are reported in the figure.

#### Supplementary Figure 9

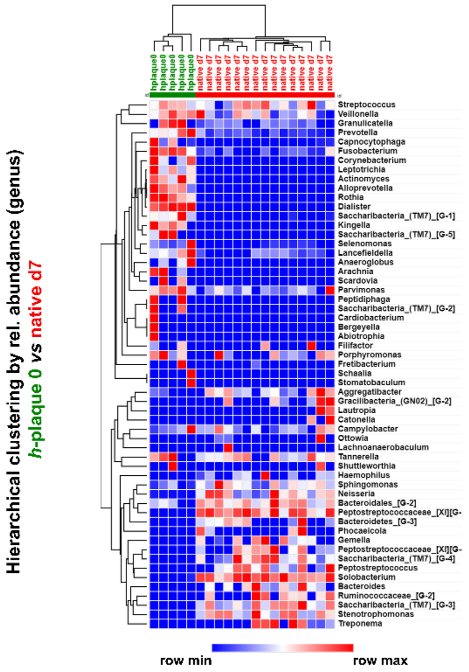


**Supplementary Figure 9. Hierarchical clustering of human subgingival microbiome analysis within the gingival tissue model: *h*-plaque 0 *versus* native d7.** Hierarchical clustering: the dendrogram across the columns has been computed as the average linkage clustering (1-Spearman rank correlation) based on microbial composition; the dendrogram across the rows has been computed as the average linkage clustering (1-Spearman rank correlation) based on the relative abundance (genus) among the samples. Graph was generated using open-source software https://software.broadinstitute.org/morpheus/

#### Supplementary Figure 10

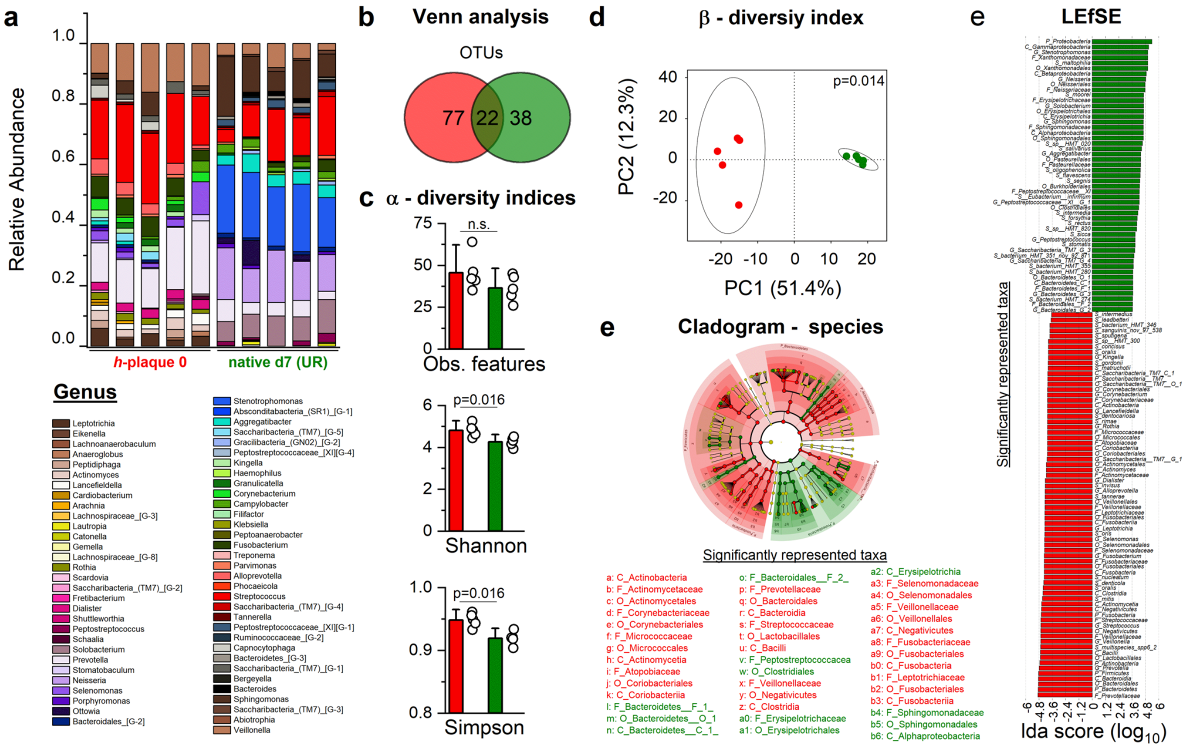


**Supplementary Figure 10. Phylogenetic studies of human subgingival microbiome distribution within the gingival tissue model: original *versus* upper regions.** Comparison between the original sample (red, $N_{patients}$=8) and the upper region (green, n=5) isolated from the gingival tissue model cultured in the gingival bioreactor (*h*-plaque 0 *vs* native day 7-UR) of: **a)** Relative Abundance – Genus; **b)** Venn analysis - species; **c)** Alpha-diversity indices: Observed features, Shannon and Simpson, Kruskal-Wallis H test, p-values (<0.05) are shown in the figure; **d)** Beta-diversity index plotted using Principal Component Analysis (PCA) by means of Euclidean distance (Aitchison distance), PERMANOVA test, p-value (<0.05) is shown in the figure; **e-f)** Differential abundance analysis showing statistically significant taxonomic levels by means of a cladogram-species (red and green nodes indicate statistically significant species, while yellow is not significant; the diameter of each circle is proportional to the abundance of the taxon represented) and taxa associated with statistically significant enriched bacteria between two groups by linear discriminant analysis Effect Size (LEfSE). Names of statistically significant taxa are reported in the figure.

### Supplementary Figure 11

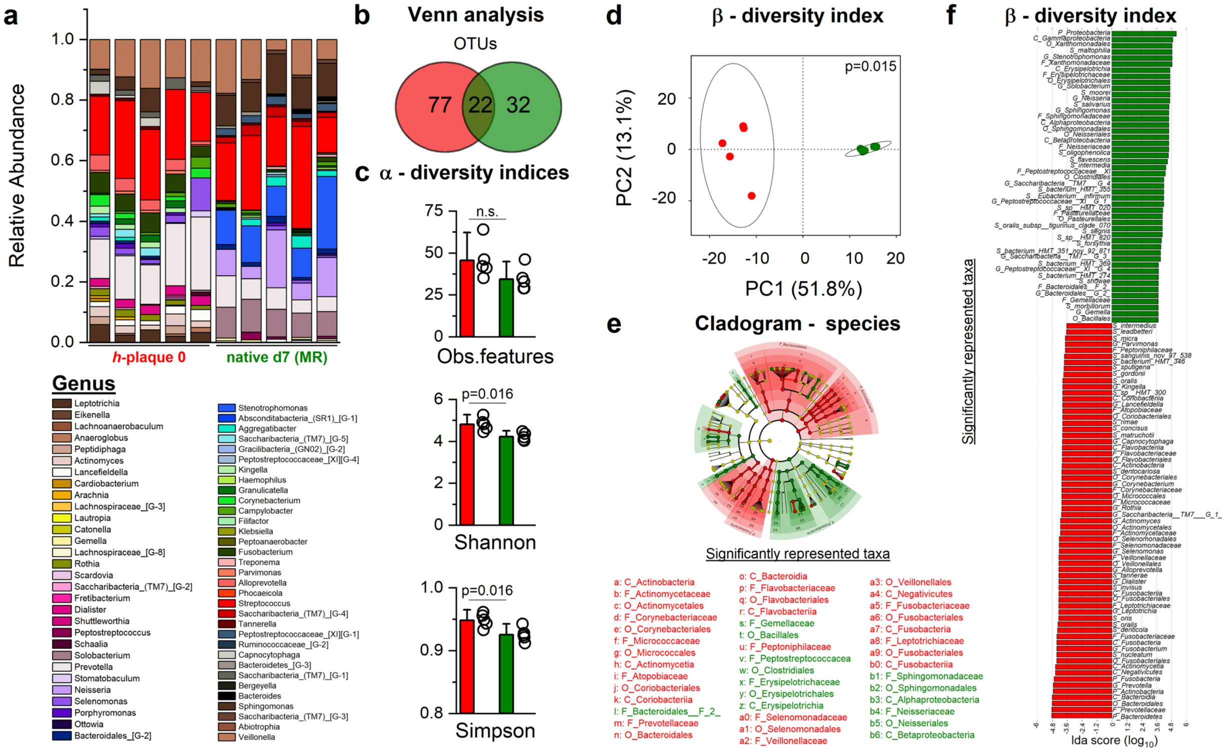


**Supplementary Figure 11. Phylogenetic studies of human subgingival microbiome distribution within the gingival tissue model: original *versus* medium regions.** Comparison between the original sample (red, $N_{patients}$=8) and the medium region (green, n=5) isolated from the gingival tissue model cultured in the gingival bioreactor (*h*-plaque 0 *vs* native day 7-MR) of: **a)** Relative Abundance – Genus; **b)** Venn analysis - species; **c)** Alpha-diversity indices: Observed features, Shannon and Simpson, Kruskal-Wallis H test, p-values (<0.05) are shown in the figure; **d)** Beta-diversity index plotted using Principal Component Analysis (PCA) by means of Euclidean distance (Aitchison distance), PERMANOVA test, p-value (<0.05) is shown in the figure; **e-f)** Differential abundance analysis showing statistically significant taxonomic levels by means of a cladogram-species (red and green nodes indicate statistically significant species, while yellow is not significant; the diameter of each circle is proportional to the abundance of the taxon represented) and taxa associated with statistically significant enriched bacteria between two groups by linear discriminant analysis Effect Size (LEfSE). Names of statistically significant taxa are reported in the figure.

#### Supplementary Figure 12

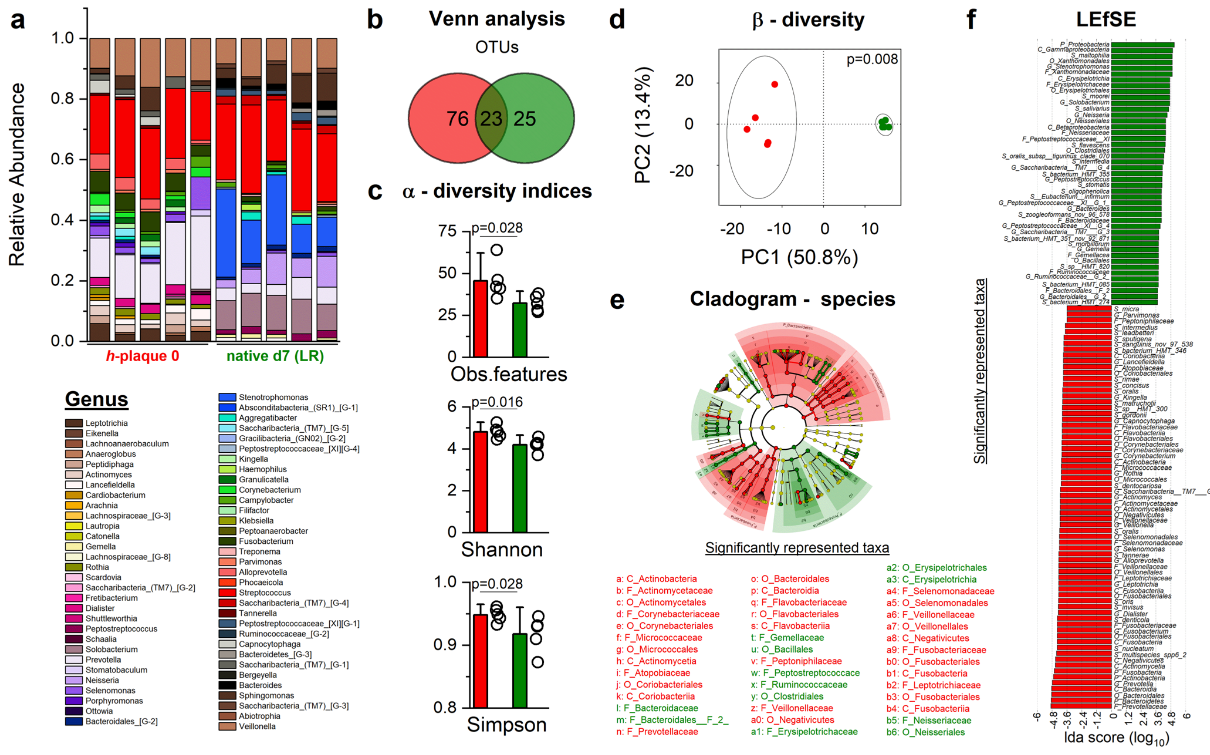

**Supplementary Figure 12. Phylogenetic studies of of human subgingival microbiome distribution within the gingival tissue model: original *versus* lower regions.** Comparison between the original sample (red, $N_{patients}$=8) and the lower region (green, n=5) isolated from the gingival tissue model cultured in the gingival bioreactor (*h*-plaque 0 *vs* native day 7-LR) of: **a)** Relative Abundance – Genus; **b)** Venn analysis - species; **c)** Alpha-diversity indices: Observed features, Shannon and Simpson, Kruskal-Wallis H test, p-values (<0.05) are shown in the figure; **d)** Beta-diversity index plotted using Principal Component Analysis (PCA) by means of Euclidean distance (Aitchison distance), PERMANOVA test, p-value (<0.05) is shown in the figure; **e-f)** Differential abundance analysis showing statistically significant taxonomic levels by means of a cladogram-species (red and green nodes indicate statistically significant species, while yellow is not significant; the diameter of each circle is proportional to the abundance of the taxon represented) and taxa associated with statistically significant enriched bacteria between two groups by linear discriminant analysis Effect Size (LEfSE). Names of statistically significant taxa are reported in the figure.

#### Supplementary Tables

##### Supplementary Table 1

| ***Cohen's effect size (d):*** *α-diversity indices* | | | |
|---|---|---|---|
| Comparison: | Observed features | Shannon | Simpson |
| Day 0 *vs* Day 0 | 4.226 | 3.344 | 1.438527786 |
| Day 0 *vs* Day 7 UR | 0.962836263 | 1.996570288 | 2.717011104 |
| Day 0 *vs* Day 7 MR | 1.214453424 | 2.291428284 | 2.05869681 |
| Day 0 *vs* Day 7 LR | 1.576470588 | 2.011351077 | 1.417956649 |
| Day 0 *vs* Day 7 (all regions) | 1.249777957 | 2.137965573 | 1.881025239 |
| Day 7: UR *vs* MR | 0.269557473 | 0.206016172 | -0.5681569 |
| Day 7: UR *vs* LR | 0.646153865 | 0.14033766 | 0.344522452 |
| Day 7: MR *vs* LR | 0.366666602 | 0.29044994 | 0.055872569 |
| Static - Day 0 *vs* Day 3 UR | 7.16138871 | 13.19437633 | 17.58223607 |
| Static - Day 0 *vs* Day 3 MR | 6.235385773 | 8.851862522 | 4.623567596 |
| Static - Day 0 *vs* Day 3 LR | 6.44831488 | 10.50931108 | 12.03260984 |
| Native - Day 0 *vs* Day 3 UR | 6.524059885 | 10.83698352 | 6.233087816 |
| Native - Day 0 *vs* Day 3 MR | 6.278900319 | 12.20083069 | 12.98470207 |
| Native - Day 0 *vs* Day 3 LR | 6.603304225 | 15.05609621 | 7.853485807 |
| Native - Day 0 *vs* Day 3 (all regions) | 6.460770846 | 12.3050555 | 7.530634328 |
| Native - Day 3 *vs* Day 7 (all regions) | 2.015998659 | 3.991987647 | 2.737029244 |

Cohen's effect size computed for all the alpha diversity indices for Figures 4,5,6,7 and Supplementary Figures 4,10,11,12 by using the effect size calculator https://lbecker.uccs.edu/.

##### Supplementary Table 2

| Time point | p-value |
|---|---|
| Day 1 – Day 0 | *<0.0001 |
| Day 1 – Day 2 | *0.00169 |
| Day 1 – Day 3 | *0.01563 |
| Day 1 – Day 4 | *0.00282 |
| Day 1 – Day 5 | *$1.07484 \times 10^{-4}$ |
| Day 1 – Day 6 | *<0.0001 |
| Day 1 – Day 7 | *$2.19647 \times 10^{-4}$ |

Statistical analysis of Supplementary Figure 6D (left). One-way ANOVA repeated measures with Dunnet's post-hoc (relative to d1); $p<0.05$.Red asterisks indicate statistically significant p-values.

**Supplementary Table 3**

| Time point | p-value |
| --- | --- |
| Day 0 – Day 3 | 0.00175 |
| Day 0 – Day 7 | 0.04288 |
| Day 1 – Day 2 | 0.00588 |
| Day 2 – Day 3 | <0.0001 |
| Day 2 – Day 4 | 0.00891 |
| Day 2 – Day 5 | 0.00775 |
| Day 2 – Day 7 | $3.83922 \times 10^{-4}$ |
| Day 3 – Day 6 | 0.00341 |

Statistical analysis of the statistically significant p-values of Supplementary Figure 6E. One-way ANOVA repeated measures with Bonferroni's post-hoc; $p<0.05$.
